# Highly heterozygous *Citrus changshan-huyou* Y. B. Chang originated from ancient hybridization between mandarin and pummelo and displayed distinct tissue-specific allelic imbalance

**DOI:** 10.1101/2025.03.24.644872

**Authors:** Zhanghui Zeng, Yingjie Luo, Haifei Hu, Lan Lan, Baojin Guo, Ping Zhou, Cong Tan, Xiaoping Huang, Tuo Qi, Zhehao Chen, Zhiming Yu, Lilin Wang, Taihe Xiang, Chengdao Li, Yong Jia

**Author notes:** Correspondence: Dr. Zhanghui Zeng, Dr. Yong Jia, Prof. Chengdao Li.

## Abstract

The genus Citrus is characterized by a reticulate evolutionary history with frequent hybridization, making it an intriguing subject for genome evolution investigation. *Citrus changshan* Y. B. Chang (Huyou) is a unique landrace first discovered in Zhejiang Province, China with premium fruit quality. The evolutionary origin of Huyou has puzzled local botanists and growers. Here, we sequenced a 120-years-old “ancestral tree” of Huyou using PacBio long read and Hi-C sequencing and assembled 2 high-quality haplotype-resolved genomes HY1 and HY2. Huyou displayed a genome heterozygosity level at 3.07%, among the highest in published citrus genomes. Using a k-mer-based tracing approach, we explicitly resolved that HY1 genome contained 87.8% mandarin, 7.3% pummelo, 0.2% citron origin, whereas HY2 had 85.0% pummelo, 2.9% mandarin, 0.3% citron, implying a hybridization event between mandarin and pummelo. Phylogeny dating showed that HY1 (2.0 Mya) and HY2 (2.18 Mya) had diverged earlier than the split of *Citrus clementina* and *Citrus reticulata*, and the split of *Citrus grandis* and *Citrus maxima*, respectively. We observed clear chromosomal recombination on chr8 and chr9 in HY1, which may have occurred after the ancestral hybridization. Further transcriptome analyses in 6 tissues revealed a strong allelic dominance of HY2 over HY1 in root tissue and moderately in stem, leaf, flower, and fruits. KEGG enrichment analyses revealed that genes related to antioxidants biosynthesis and lipid metabolisms were most significantly affected by allelic imbalance. This first report of allelic imbalance in citrus species support Huyou as an interesting model to investigate genome evolution following distant hybridization.

## Introduction

The genus Citrus belongs to the Aurantioideae subfamily of Rutaceae and contains a diverse array of species such as sweet orange (*C. sinensis*), mandarin (*C. reticulata*), pummelo (*C. grandis*), citron (*C. medica*), and lemon (*C. limon*). Their diverse flavors, nutritional benefits, and industrial applications have made them beloved fruits worldwide. Citrus ranks the third largest fruits in term of production volume (https://www.statista.com/) and hold substantial agricultural, economic, and cultural values globally (Gmitter*, et al.*, 2012). Beyond their economic and cultural importance, citrus fruits present a fascinating subject for genomic and evolutionary studies due to their intricate origins and diversification. The genus Citrus is characterized by a reticulate evolutionary history, marked by frequent hybridization and polyploidization events (Wu*, et al.*, 2018). This complex evolutionary past has led to various cultivated species or landraces with different heterozygosity levels, making citrus an intriguing subject for genome evolution investigation.

*Citrus changshan-huyou* Y. B. Chang (Huyou) is a unique Citrus species originated in the Changshan County of the Zhejiang Province in China (Li*, et al.*, 2019; Yin-bin, 1991). The earliest cultivation of Huyou has been recorded over 600 years ago, well before its massive planting and production over 100 years as a prominent landrace. Its fruit has a golden color and contains abundant components, such as amino acids, vitamins, naringin, limonin, citrate and other nutrients (Zhang*, et al.*, 2012; Guo*, et al.*, 2018; Sheng*, et al.*, 2017). These components not only invested the fruit unique aroma and taste that combines sweet, sour, and a bit of bitterness, but also health benefits, including lowering blood sugar, hepatoprotective and anti-inflammatory effects (Zhang*, et al.*, 2012; Guo*, et al.*, 2018; Jiang*, et al.*, 2019). These benefits have led to the widespread consumption of Huyou in Zhejiang and neighboring province. The economic importance and scientific values of Huyou are attracting increasing attention. A recent scientific dataset reported the haplotype-resolved genome assembly of a Huyou accession and transcriptome sequencing in 6 tissues (Miao*, et al.*, 2024). Despite the high quality Huyou genomic data, the dataset fell short on genomic origin and evolution analysis for this important natural hybrid citrus landrace. The domestication and evolutionary origin of Huyou remains to be characterized.

Over the past decade, numerous citrus species have been sequenced, generating genomic data that has significantly advanced our knowledge of citrus biology and genome evolution. Key milestones include the sequencing and assembly of reference genomes for important citrus species such as sweet orange (*C. sinensis*) (Xu*, et al.*, 2013), clementine mandarin (*C. clementina*) (Wu*, et al.*, 2014), and pummelo (*C. maxima*) (Wu*, et al.*, 2014). These were followed by whole genome resequencing for large citrus germplasm collections that encompass diverse citrus wild, landraces, and cultivars lines (Wu*, et al.*, 2018; Wang*, et al.*, 2017). All these studies have highlighted a complex history of admixture and hybridization during citrus domestication. The sequencing projects indicate that modern citrus varieties are often the result of crosses between a few ancestral species such as mandarin, pummelo, and citron. Beyond the genomic insights into genome origin and evolution, citrus genome sequencing studies have also led to the identification of genes associated with important agronomic traits, including fruit quality, disease resistance, and stress tolerance. For example, studies have identified genes involved in flavor synthesis in lemon (Bao*, et al.*, 2023), disease resistance in trifoliate orange (Peng*, et al.*, 2020), and citric acid accumulation in oranges (Huang*, et al.*, 2023). Recently, citrus pangenome data for various citrus and citrus related species have been made available (Huang*, et al.*, 2023; Liu*, et al.*, 2022). These genomic resources allow researchers to delve deeper into the genetic basis of key traits, paving the way for more targeted and efficient breeding strategies (Liu*, et al.*, 2022; Humann*, et al.*, 2022).

Haplotype-phased genome refers to a genome sequence where the two sets of chromosomes inherited from each parent are separately distinguished and assembled. The assembly of highly accurate haplotype-phased genomes is achieved through advanced sequencing technologies such as long-read sequencing (e.g., PacBio HiFi, Nanopore), Hi-C sequencing (chromosome conformation capture). It allows for the identification of allele-specific variations, structural variations, and gene expression patterns that would be obscured in a standard genome assembly. This is particularly crucial for genomic study on heterozygous organisms such as Citrus species, which are marked with frequent hybridization events. Recently, several haplotype-resolved genome assemblies have been generated, including *C. limon* (Mario*, et al.*, 2021), *C. australis* (Nakandala*, et al.*, 2023), *C. changshanensis* (Miao*, et al.*, 2024), *Papeda* (Wang*, et al.*, 2025), and a haploid genome assembly for pummelo (Wang*, et al.*, 2017), offering a more detailed view of their genetic makeup.

In this study, we performed genome sequencing and assembly of a 120-years-old “ancestor tree” of Huyou currently still growing in Changshan, Zhejiang Province where Huyou was first discovered. We integrate high coverage Illumina short read and PacBio long read sequencing with Hi-C sequencing technology for the assembly of 2 haplotype-resolved chromosome scale genomes. This is followed by in-depth genome tracing, structural variation, and comparative genomic analysis with previously established citrus ancestor species to fully characterize the unique genomic features of Huyou and to resolve its evolutionary origin. Furthermore, assisted by previously published transcriptome data and the haplotype-resolved genome assemblies in this study, we comprehensively characterized the allelic imbalance across different tissues. Gene family expansion/contraction and transcriptome profiles for key genes from important metabolic pathways related to citrus fruit quality were also analyzed. We highlight the exceptional genome heterozygosity of Huyou compared to known Citrus genomes and the unique tissue-specific allelic imbalance in the root tissue. Our study supports Huyou as a novel genomic resource for citrus breeding and also an interesting model to investigate genome evolution following distant hybridization.

## Materials and Methods

### Plant materials and nucleic acid extraction

Fresh leaves, shoots and fruits were collected from a 120-year-old *C. changshan-huyou* Y. B. Chang tree in Chengtan Village, Qingshi Town, Changshan County (118 30′E, 28 51′N), Zhejiang Province, China, in the spring of 2022 **(Figure 1A)**. These tissues were quickly frozen in liquid nitrogen and then stored at −80℃ before DNA and RNA extraction. Total genomic DNA was extracted from the leaf tissue, and its quality was assessed by the Qubit 3.0 Fluorometer (Thermo Fisher, USA), while the total RNA from leaves, shoots and fruits was extracted using TRIzol Reagent (Invitrogen). The quantity and quality of the RNA were examined using NanoDrop One spectrophotometer (Thermo Fisher, USA) and an Agilent 2100 Bioanalyser (Agilent Technologies), respectively.

**Figure 1.**
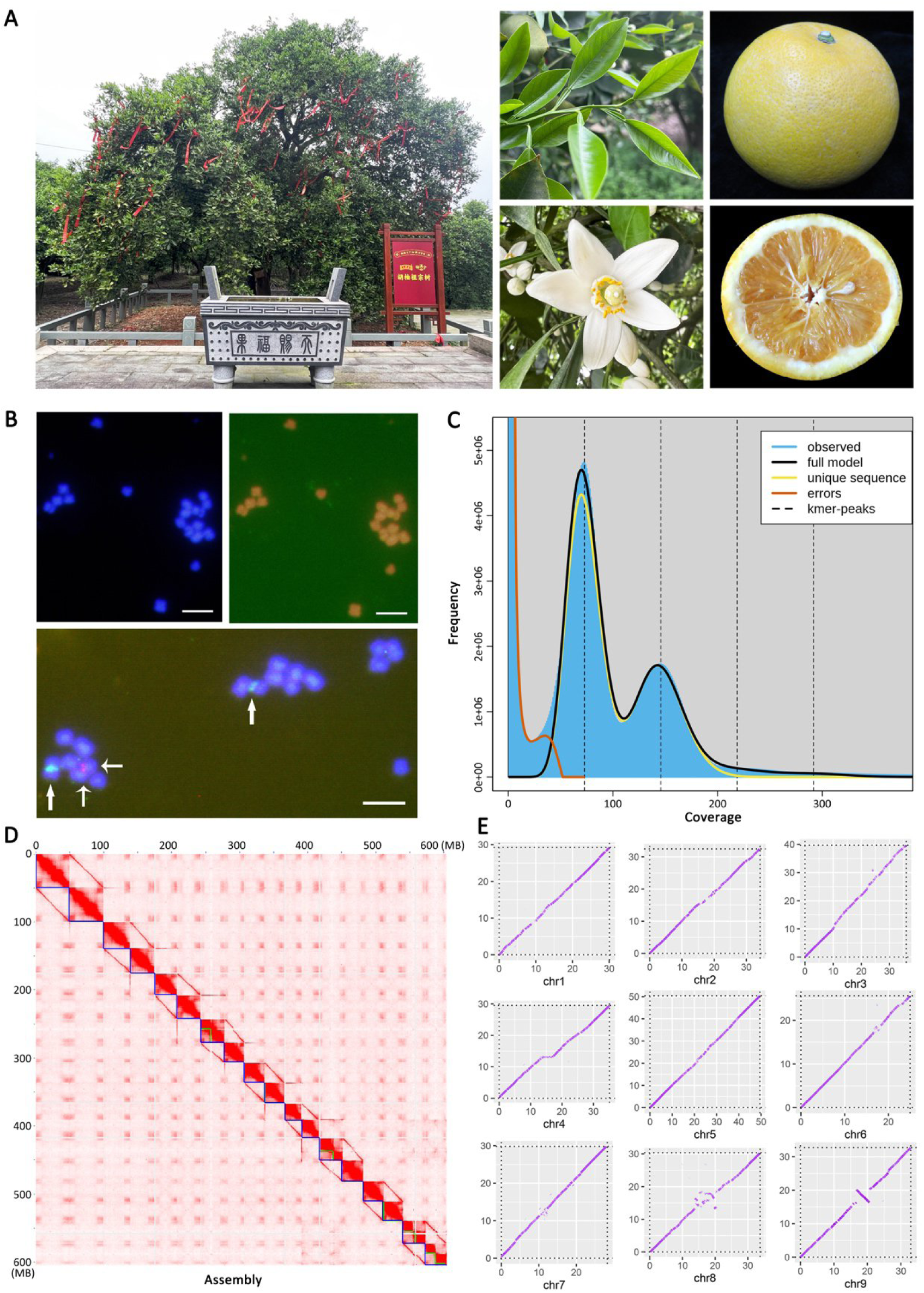
Huyou tree characteristics, karyotype analysis, and Genome survey. **A.** The 120 years old “ancestral” Huyou tree sequenced in this study (whole tree, stem, leaves, flowers, and fruits). **B**. Karyotype analysis on chromosomes of Huyou root tips. Karyotype pictures of Huyou using DAPI staining (top left), FISH analysis with a telomere specific repeat probe (top right), and 5S rDNA (red) and 18S rDNA (green) repeat sequence probes (bottom). Bar = 5 μm. **C**. Genome size and heterozygosity estimation based on K-mer frequency analysis (K = 21). The fist peak reveals heterozygous k-mers while the second peak indicates the homozygous k-mers. **D**. Hi-C interaction heatmap for the assembled HY1 and HY2 genomes. Blue lines indicated chromosomal boundary. The 18 chromosomes were sorted based on length in decreasing order. The bottom-right cluster represented unanchored contigs. **E**. Dot plots displaying the synteny between HY1 and HY2 for each chromosome.

### Genome karyotype identification

Karyotype analysis was conducted with fluorescence *in situ* hybridization (FISH) method as previously described (Wang*, et al.*, 2017; Zhang*, et al.*, 2022), and was performed by OMIX Technologies Corporation (Chengdu, China). Briefly, excised young root tips (2-3cm) of Huyou via tissue culture (Li*, et al.*, 2017) were treated with nitrous oxide gas for 2h under 1 MPa, then fixed in ice-cold 90% acetic acid for 10 min. After washing in water, the root tips were diced and digested in 1% pectolyase Y23 and 2% cellulase Onozuka R-10 (Yakult Pharmaceutical, Japan) for 1 h at 37℃. After digestion, the root sections were washed in 70% ethanol three times briefly, then broken by using a needle and vortexed at maximum speed in 100% ethanol for 30 sec at room temperature to separate cells from one another. The cells were collected at the bottom of the tube by centrifugation (4000r/min) and resuspended in glacial acetic acid. The cell suspension was dropped onto glass slides in a box lined with wet paper. A telomere-specific repeat probe 5′-(TTTAGGG)_6_-3′) was used to detect the number of intact chromosomes, and 5s rDNA and 18s rDNA repeat sequence probe were used to identify multiple copies of chromosomes. The fluorescence staining of the chromosomes was performed using 4′,6-diamidino-2-phenylindole (DAPI). After DAPI staining, the dispersed metaphase chromosome cells were counted under a fluorescence microscope (Zeiss LSM880, Germany).

### Genome size and heterozygosity estimation

For genome profiling, the DNA library was constructed with insert sizes of ∼300 bp using Nextera DNA Flex Library Prep Kit (Illumina, San Diego, CA, USA), and subsequently sequenced on the Illumina NovaSeq 6000 platform (Illumina, San Diego, CA, USA), which generated 150 bp pair-end reads. The software FastQC (version: 0.20.1) was used to filter the original reads and discard the low-quality reads. A total of 57.05 Gb clean data were generated by Illumina sequencing and were used to perform the genome survey for Huyou. Reference-free estimation of genome size and heterozygosity were performed using a k-mer frequency approach implemented in GenomeScope 2.0 (Ranallo-Benavidez*, et al.*, 2020).

### PacBio, Hi-C and RNA Sequencing

To generate long-read sequencing reads for Huyou, DNA libraries were prepared using the SMRTbell Express Template Preparation kit 1.0 following the PacBio 20-kb protocol (https://www.pacb.com/), and sequenced on the PacBio Sequel II platform (Pacific Biosciences of California, Inc.). After the data quality control, by removing sequencing adaptors and filtering low-quality short read, and correcting errors via SMRT link (v8.0), 26.17 Gb of HIFI reads were obtained (**Table 1**).

**Table 1.** The statistics for genome assembly and gene annotation of Huyou. HY1 and HY2 refers the two haplotype genomes.

| Sequencing platform | Illumina, PacBio, Hi-C |  |
| --- | --- | --- |
| **Assembly** | HY1 | HY2 |
| Chromosome number | 9 | 9 |
| Total genome size | 326.65 Mb | 322.79 Mb |
| Size anchored to chromosome | 299.73 Mb / 91.8% | 304.81 Mb / 94.4% |
| Number of contigs | 324 | 235 |
| N50 contigs | 21.15 Mb | 18.77 Mb |
| Largest contig | 46.45 Mb | 34.26 Mb |
| Complete BUSCOs | 98.1% | 97.7% |
| GC content | 56.60% | 55.02% |
| **Genome annotation** |  |  |
| Number of protein-coding genes | 42091 | 42050 |
| Mean mRNA length | 2570 bp | 2554 bp |
| Mean CDS length | 1245 bp | 1245 bp |
| Mean exons per gene | 5.75 | 5.77 |
| Mean exon length | 216.40 | 215.87 |
| Number of TEs |  |  |
| DNA | 14097 | 15545 |
| LINE | 6762 | 7428 |
| SINE | 248 | 269 |
| LTR | 50487 | 56301 |
| Total size of TEs | 181.5 Mb / 55.56% | 179.4Mb / 55.58% |
| Number of non-coding RNA |  |  |
| miRNA | 161 | 150 |
| tRNA | 1003 | 790 |
| rRNA | 1590 | 2686 |
| snoRNA | 1393 | 1463 |

The generation of a Hi-C library was performed as described previously (Strijk*, et al.*, 2021). Briefly, the genomic DNA was random broken into 300-700 bp fragments via DNA cross-linking, restriction enzyme digestion, cohesive end repair, DNA cyclization, and DNA purification. Then the sample was sequenced on an Illumina NovaSeq 6000 platform (PE 150). After the removal of sequencing adaptors and low-quality reads using fastp software (v 0.21.0) (Chen*, et al.*, 2018), 31.77 Gb Hi-C clean reads were obtained.

RNA sequencing (RNA-seq) samples were obtained by mixing equal amounts of RNA extracted from each tissue (leaf, shoot, and fruit) and used for cDNA library construction. After sequencing on the Illumina NovaSeq 6000 platform, approximately 10 Gb of sequencing data was obtained.

### De novo genome assembly and annotation

Haplotype-resolved *de novo* genome assembling was performed using hifiasm software (Cheng*, et al.*, 2021) with HIFI reads and Hi-C phasing. The default k-mer size was changed 19 to obtain correct estimation heterozygous site coverage. The resulted haplotype-resolved genome contigs were further purged for duplicates using Purge_Dups (Guan*, et al.*, 2020) by aligning the HIFI reads back to the assembled genomes using minimap2 tool (Li, 2018). The purged haplotype genomes were scaffolded into chromosome groups using the pipeline implemented at https://github.com/WarrenLab/hic-scaffolding-nf, which employs Chromap (Zhang*, et al.*, 2021) and YaHS (Zhou*, et al.*, 2023) for sequence alignment and Hi-C scaffolding, respectively. Genome statistics were calculated using the assembly-stats tool (https://github.com/sanger-pathogens/assembly-stats). Then, the juciebox software (v1.11.08) (Durand*, et al.*, 2016) was used to draw Hi-C heatmap based on the interaction intensity and relative position between contigs. Following manual correction, the uniquely mapped and valid interaction paired-end reads were used to build the pseudochromosome sequences. The pseudochromosomes were sorted and oriented using C. sinensis as the reference by RagTag (Alonge*, et al.*, 2022).

The quality of the assembled genomes was evaluated by Benchmarking Universal Single-Copy Orthologs against the eukaryota_odb10 database (BUSCO v.4.1.4) (Simao*, et al.*, 2015). Repeat sequences of the haplotype genomes were searched using an automated pipeline implemented by EarlGrey (Baril*, et al.*, 2024). The soft-masked genomes were used for gene model prediction using GeMoMa v1.8 (Keilwagen*, et al.*, 2019) based on homology and transcriptome evidence. For homology references, the gene models from three ancestor genomes (C. sinensis, C. ichangensis, C. medica) were used. For transcriptome evidence, the RNA sequencing reads were mapped to the masked genomes using STAR (Dobin*, et al.*, 2013) and the resulted alignment files were used as input for GeMoMa. For noncoding RNA annotation, the transfer RNAs (tRNAs) were annotated by tRNAscan-SE (v1.23) (Chan*, et al.*, 2021). Other non-coding ribosomal RNAs (rRNAs), small nuclear RNAs (snRNAs) and microRNAs (miRNAs) were identified using the Infernal toolkits (Nawrocki*, et al.*, 2009) based on information from the Rfam database. Gene function annotations were performed using Interproscan for GO annotation and kofam_scan for KEGG annotation. GO and KEGG enrichment analyses for Huyou genome specific genes were performed using TBtools (Chen*, et al.*, 2020). The circos v0.69 (Krzywinski*, et al.*, 2009) was used to visualize genomic characteristic information, including gene density, GC density, TE density, gene density, and synteny block.

### Variants, synteny, and genome origin analyses

The genomic variations and synteny across the haplotype genomes of Huyou and their ancestor genomes *C. reticulata* (mandarin, NCBI accession ID: ASM325862v1) (Wang*, et al.*, 2018) and *C. maxima* (pummelo, NCBI accession ID: ASM2964120v1) (Zheng*, et al.*, 2023) were analyzed using SyRI software (Goel*, et al.*, 2019). Genome alignment was performed using minimap2 (Li, 2018). The ancestral origin of Huyou haplotype genomes were determined using the k-part tool at https://github.com/sc-zhang/kPart. Variations between HY1 and HY2 were counted per 1 Mb windown based on the output from SyRI. For ancestor tracing, genomic data for 3 ancestor genomes (*C. reticulata*, *C. maxima*, *C. medica*) were downloaded and used as references. The ancestor of Huyou genomes were assigned for each 100 kb window using kPart with -k 21 parameter (https://github.com/sc-zhang/kPart). The percentages of each ancestor origin were calculated. The haplotype origin for each chromosome was also phased and classified using SubPhaser program with a similar k-mer based method (Jia*, et al.*, 2022).

### Orthologs inferences, phylogeny development, and gene family analyses

Protein sequences of Huyou genomes and 19 other species, including *Aquilegia coerulea*, *Mangifera indica*, *Vitis vinifera*, *Malus domestica Golden*, *Solanum lycopersicum*, *C. sinensis*, *C. reticulata*, *C. maxima*, *C. medica*, *C. clementina*, *Poncirus trifoliata*, *Fortunella hindsii*, *C. ichangensis*, *Atalantia buxfoliata*, and *Litchi chinensis Sonn* (downloaded from Citrus Genome Database at https://www.citrusgenomedb.org/) (Liu*, et al.*, 2022) were used as input for OrthoFinder v2.3.12 (https://github.com/david emms/Ortho Finder) (Emms and Kelly, 2019) for orthologs inferences. For alternative transcripts, only the sequences for the longest transcripts were used. The obtained species tree was used as input for downstream Bayesian estimation of divergence time using mcmctree from PAML v4.9 toolkits (Yang, 2007). The CDS sequences for the identified single copy genes from Orthofinder were extracted. Codon-based sequence alignments were performed using a combination of muscle and seqmagick backtrans-align functions. The concatenated CDS alignment was trimmed using Gblocks before input into mcmctree. Five divergence time points derived from https://timetree.org/ were used to calibrate the species tree. The final obtained species tree from mcmctree and orthologous gene count results from Orthofinder were input into café for gene family expansion/contraction analyses. The gene family changes along the species tree were visualized using tvBOT (Xie*, et al.*, 2023) online tool at https://www.chiplot.online/tvbot.html. For gene copy number analysis, the gene counts for corresponding hierarchical orthologoups from Orthofinder outputs were used.

### Genome divergence time analyses

The genome divergence time was assessed by *Ks* profile analyses using WGDI program (Sun*, et al.*, 2022). Each citrus genome was compared to itself following the WGDI instruction. Protein sequences for the primary transcripts of each species were used for blastp using diamond program (Buchfink*, et al.*, 2021). Syntenic blocks were identified using the build-in function in WGDI. Codon-based sequence alignments were performed using muscle (Edgar, 2004) and pal2nal (Suyama*, et al.*, 2006) programs. Ks values calculation for collinear genes using yn00 from PAML v4.9 program (Yang, 2007). Ks peaks at p-value ≤ 0.05 were identified and fitted using WGDI. The whole genome duplication event in Vitis vinifera (Jaillon*, et al.*, 2007) was used a reference.

### Transcriptome analyses

Transcriptome RNA sequencing data in 6 Huyou tissues (root: SRR28430799, stem: SRR28430800, leaf: SRR28430798, flower: SRR28430801, unripe fruit: SRR28430803, ripe fruit: SRR28430802) were downloaded from a previous study (Miao*, et al.*, 2024). Raw reads were trimmed using fastp program with --qualified_quality_phred 20 and --length_required 30 parameters. The resulted clean reads were mapped to merged haplotype genomes using STAR program, then were quantified using RSEM (Li and Dewey, 2011) based on the annotated gene models in this study. The syntenic gene pairs between HY1 and HY2 were identified using MCScanX program (Wang*, et al.*, 2012). The comparison of syntenic genes expression was performed by calculating the log2 value of the ratio of HY1 gene expression to HY2 gene expression.

## Results

### Genome assembly and annotation of the 120 years-old “ancestral tree” of *Citrus changshan-huyou* Y. B. Chang

In this study, a century-old Huyou tree (120 years old as of 2025) currently growing and protected in Chentang Village, Changshan County, after which Huyou was named by Mr. Yunbing Chang in 1991, was selected for genome sequencing (**Figure 1A**). This century-old tree was believed by many local Huyou growers and botanists to be the ancestor of most Huyou trees currently cultivated across Zhejiang province and was thus termed as the “ancestral tree”. Notably, this “ancestral tree” of Huyou, despite its old age, remains highly vigorous and productive with premium fruits to date (**Figure 1A**, photographed in 2024).

To determine the chromosome number and ploidy level of Huyou, high-resolution metaphase chromosome preparation assay was performed using root tip cells from tissue cultured Huyou seedlings. Results showed that the genome of Huyou is composed of 18 chromosomes (2n = 2x = 18, **Figure 1B**), which was confirmed by FISH with 5S rDNA and 18S rDNA repeat sequence probes, and consistent with those reported in other citrus species (Wang*, et al.*, 2017). *K*-mer frequency distribution analysis (*K* = 21) of Illumina short reads (57.05 Gb, ∼189 × coverage) showed a distinct bimodal profile with 3.07% heterozygosity (**Figure 1C**), among the highest across all citrus genomes reported to date (Wu*, et al.*, 2018; Miao*, et al.*, 2024).

Due to its exceptionally high heterozygosity level, long-read PacBio HIFI sequencing (26.17 Gb, ∼ 86 ×) and the Hi-C sequencing (31.77 Gb, 105 ×) were performed to obtain high-quality chromosome-scale reference assembly for Huyou. The de novo assembling using Hifiasm with HIFI long reads and Hi-C reads as inputs produced two haplotype genomes HY1 (324 contigs, total size at 326.65 Mb with contig N50 of 21.15 Mb) and HY2 (235 contigs, total size at 322.79 Mb with contig N50 of 18.77 Mb) (**Table 1**). GC contents for HY1 (56.6%) and HY2 (55.02%) were comparable with each other. The largest contigs for HY1 and HY2 were at 46.45 Mb and 34.26 Mb, respectively. The assembled contigs were anchored into 9 pairs of chromosomes with Hi-C scaffolding. Distinct chromosomal groupings on HI-C interaction heatmap were observed in **Figure 1D**, indicating the high quality of genome anchoring. The chromosomes were further ordered and named using *C. sinensis* (v3.0 from the Citrus Genome Database http://citrus.hzau.edu.cn/download.php) as reference. Particularly, 91.8% and 94.4% of the contigs for HY1 and HY2, respectively, could be assigned to chromosomes. Using the eukaryota_odb10 database as reference, the Benchmarking Universal Single-Copy Orthologs (BUSCO) revealed completeness of 98.1% and 97.7% for HY1 and HY2, respectively. Using HY1 as the reference, genome alignment of HY1 and HY2 revealed overall well-conserved synteny between the 2 haplotype genomes, albeit some large structural variations were observed mainly in the central region of each chromosome, such as a large insertion in chr4 and large inversions in chr8 and chr9 (**Figure 1E**). Despite the overall synteny, the mosaic pattern and gaps in the genomic alignment plot indicated significant genetic variations between HY1 and HY2 (**Figure 1E**), consistent with the exceptional high heterozygosity in Huyou observed in this study.

Repetitive sequences were identified in the two haplotype genomes HY1 (181.5 Mb or 55.56%) and HY2 (179.4 Mb or 55.58%) (**Table 1**). In both genomes, LTR represented the largest TE family, followed by DNA, LINE, Rolling Circle, sequentially (**Supplementary File S0**). Despite a relatively smaller genome size and smaller total TE content, HY2 displayed higher number of TEs in all TE classes than that in HY1 (**Table 1**). The repeat sequences were masked before gene model prediction using both homology search (*C. sinensis*, *C. ichangensis*, *C. medica*) and RNA sequencing data (10 Gb), which led to the identification of 42091 and 42050 protein-coding genes for HY1 and HY2, respectively, with comparable mean length for transcripts, CDS, exon, and average exon number per gene (**Table 1**). HY1 displayed relatively higher number of micro-RNA (161 miRNA) and tRNA (1003) but relatively lower number of small nuclear RNA (1393 snoRNA) than that in HY2 (150 miRNA, 790 tRNA, 1463 snoRNA). However, the number of ribosomal RNA in HY2 (2686 rRNA) was much higher than that in HY1 (1590 rRNA). The distribution of TEs, genes, and non-coding RNAs in HY1 and HY2 were displayed in **Figure 2A**. Overall, we observed higher gene density in the distal regions of each chromosome compared to the central region, whereas TEs tend to be enriched in the central regions. In summary, we obtained 2 haplotype-resolved Huyou assemblies with high quality and completeness.

**Figure 2.**
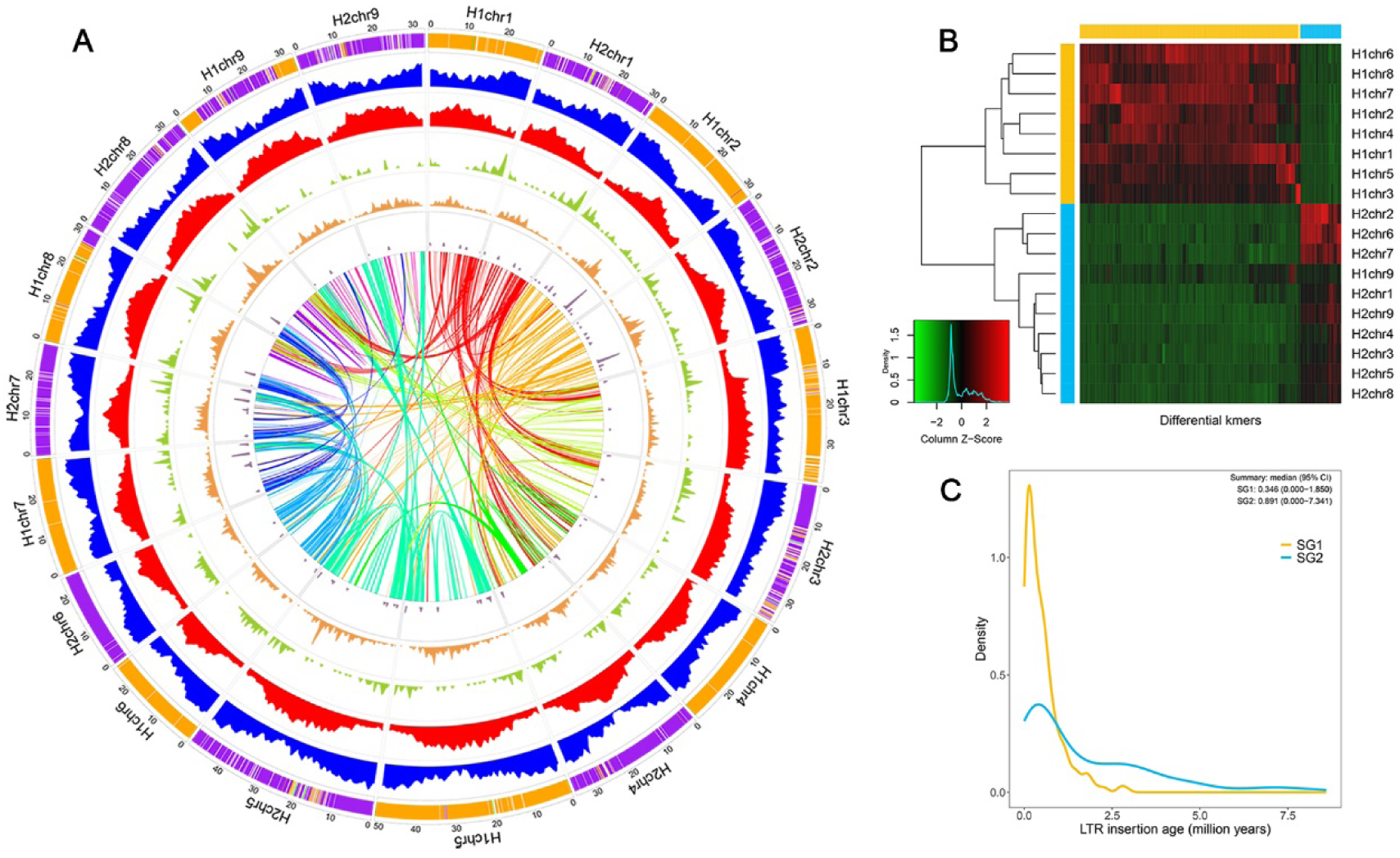
Ancestral tracing and comparative genome analysis of Huyou. **A.** Circles from outside to inside represent chromosomes of two phased haplotypes (mandarin, orange; pummelo, purple; citron, green; undetermined, white), gene density (blue), transposable element density (red), tRNA density (green), small nucleolar RNA density (orange) and microRNA density (plum). The curved lines in the center indicate patterns of gene collinearity. All distributions are drawn in a window size of 100kb. **B.** The histogram of differential k-mers among homoeologous chromosomes of HY1 and HY2. **C**. Insertion time of subgenome-specific LTR-RTs (output from SubPhaser program, see method for details).

### Ancestral tracing of Huyou haplotype genomes HY1 and HY2

Due to the high heterozygosity level and significant structural variations between the haplotype-resolved genomes, the ancestral origin of each block (100 kb) of the HY1 and HY2 genomes were determined using a k-mer based tracing approach. The distribution of differential k-mers of homologous chromosomes indicated that HY1 and HY2 could be clearly separated into 2 distinct progenitors (**Figure 2B**), except for chr9 in HY1 which displayed a mixture origin and was clustered with HY2. Using the citrus ancestor genomes mandarin (*C. reticulata*), pummelo (*C. maxima*), and citron (*C. medica*) as references, the genomic origin of HY1 and HY2 were traced and quantified. Results showed that 87.8% of HY1 genome originated from mandarin, followed by 7.3% from pummelo, 0.2% from citron, and 4.7% unknown source (could not be explicitly assigned) (**Table 2, Supplementary File S1**). In contrast, 85.0% of HY2 genome was derived from pummelo, followed by 2.9% from mandarin, 0.3% from citron, and 11.9% from unknown source (**Table 2**, **Figure 2A**). These results strongly suggested that Huyou may have originated from a hybridization between mandarin (HY1) and pummelo (HY2). At the chromosome level, chr1-chr7 of HY1 were predominantly from mandarin, whereas chr8 (11.9% pummelo) and chr9 (54.1% pummelo) of HY1 contained significant components from pummelo, which may have resulted from chromosome recombination. Mapping the genomic origin to the chromosomes indicated clear evidence of chromosome recombination in HY1 genome, whereby one distal region of chr8 and the central chr9 were replaced by pummelo genome (**Figure 2A**). For the HY2 genome, all chromosomes were predominantly from pummelo, with minor introgression of mandarin on chr1 (3.3% mandarin), chr3 (7.6%), chr5 (4.8%), chr8 (1.2%), and chr9 (3.6%) (**Table 2**). In contrast to HY1 which retained clear segmental recombination boundary, the genome introgressions in HY2 were generally mosaic along the chromosomes (**Figure 2A**), which may provide further clue on the evolution history of Huyou following the hybridization of its pummelo and mandarin parents. Furthermore, the LTR insertion age analysis also indicated distinct profiles for HY1 and HY2 (**Figure 2C**), providing additional support for two divergent progenitors. Specifically, most of the LRTs in HY1 displayed concentrated age with a single peak of insertion time, whereas the insertion time of LRTs in HY2 was relatively dispersed and contained an additional minor peak at a much older age (**Figure 2C**).

**Table 2.**
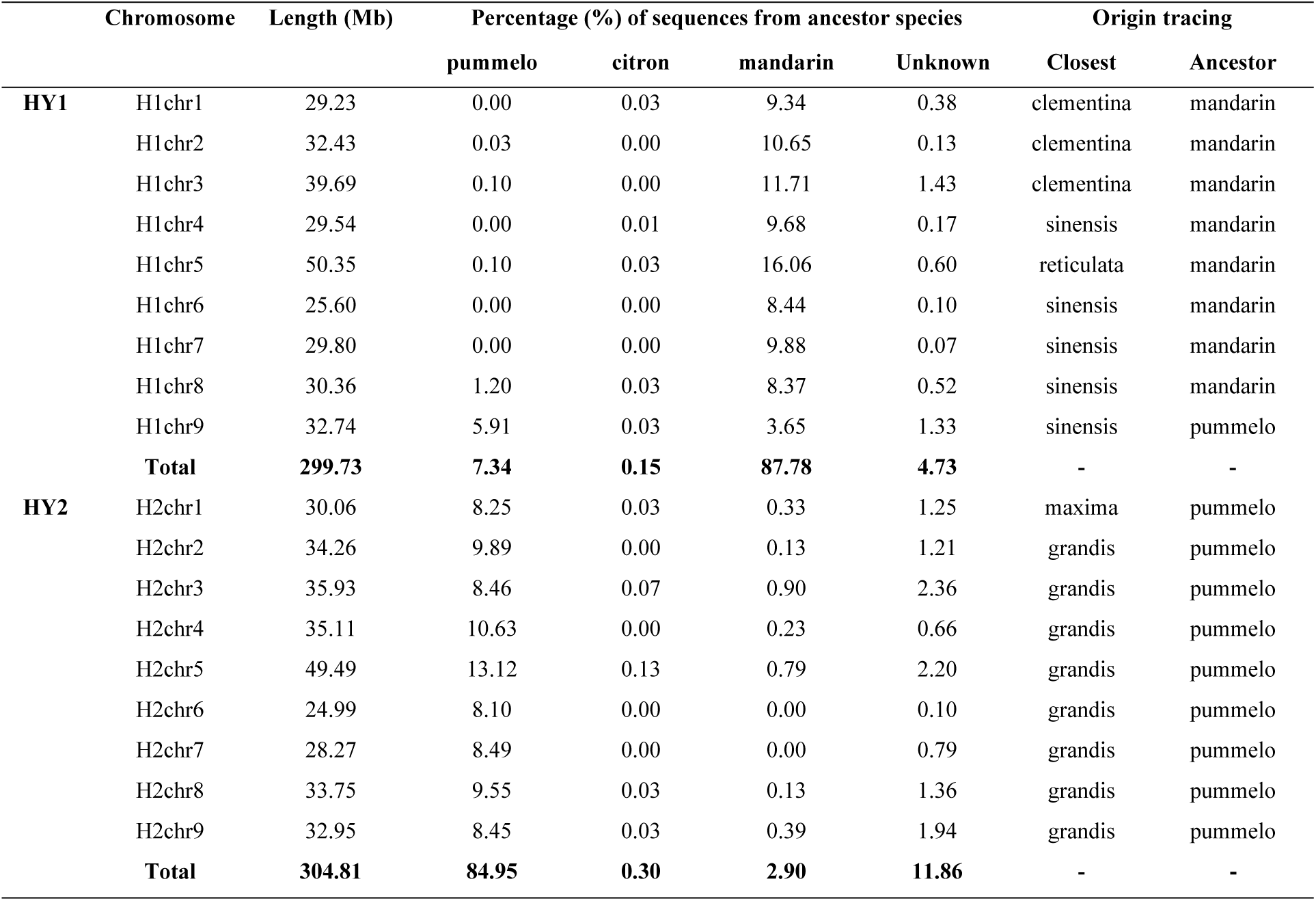
The ancestral origin of each chromosome of haplotype-resolved Huyou. Genome tracing was implemented using a k-mer-based approach by mapping sequencing data of three reference ancestral genomes mandarin (*C. reticulata*), pummelo (*C. maxima*), and citron (*C. medica*) to HY1 and HY2 using 100kb window. See the method section and **Supplementary File S1** for details.

|  | Chromosome | Length (Mb) | Percentage (%) of sequences from ancestor species |  |  |  | Origin tracing |  |
| --- | --- | --- | --- | --- | --- | --- | --- | --- |
|  |  |  | pummelo | citron | mandarin | Unknown | Closest | Ancestor |
| HY1 | H1chr1 | 29.23 | 0.00 | 0.03 | 9.34 | 0.38 | clementina | mandarin |
|  | H1chr2 | 32.43 | 0.03 | 0.00 | 10.65 | 0.13 | clementina | mandarin |
|  | H1chr3 | 39.69 | 0.10 | 0.00 | 11.71 | 1.43 | clementina | mandarin |
|  | H1chr4 | 29.54 | 0.00 | 0.01 | 9.68 | 0.17 | sinensis | mandarin |
|  | H1chr5 | 50.35 | 0.10 | 0.03 | 16.06 | 0.60 | reticulata | mandarin |
|  | H1chr6 | 25.60 | 0.00 | 0.00 | 8.44 | 0.10 | sinensis | mandarin |
|  | H1chr7 | 29.80 | 0.00 | 0.00 | 9.88 | 0.07 | sinensis | mandarin |
|  | H1chr8 | 30.36 | 1.20 | 0.03 | 8.37 | 0.52 | sinensis | mandarin |
|  | H1chr9 | 32.74 | 5.91 | 0.03 | 3.65 | 1.33 | sinensis | pummelo |
|  | **Total** | **299.73** | **7.34** | **0.15** | **87.78** | **4.73** | - | - |
| HY2 | H2chr1 | 30.06 | 8.25 | 0.03 | 0.33 | 1.25 | maxima | pummelo |
|  | H2chr2 | 34.26 | 9.89 | 0.00 | 0.13 | 1.21 | grandis | pummelo |
|  | H2chr3 | 35.93 | 8.46 | 0.07 | 0.90 | 2.36 | grandis | pummelo |
|  | H2chr4 | 35.11 | 10.63 | 0.00 | 0.23 | 0.66 | grandis | pummelo |
|  | H2chr5 | 49.49 | 13.12 | 0.13 | 0.79 | 2.20 | grandis | pummelo |
|  | H2chr6 | 24.99 | 8.10 | 0.00 | 0.00 | 0.10 | grandis | pummelo |
|  | H2chr7 | 28.27 | 8.49 | 0.00 | 0.00 | 0.79 | grandis | pummelo |
|  | H2chr8 | 33.75 | 9.55 | 0.03 | 0.13 | 1.36 | grandis | pummelo |
|  | H2chr9 | 32.95 | 8.45 | 0.03 | 0.39 | 1.94 | grandis | pummelo |
|  | **Total** | **304.81** | **84.95** | **0.30** | **2.90** | **11.86** | - | - |

### Phylogeny dating and gene family analyses of HY1 and HY2

To deduce the divergence time of HY1 and HY2, single copy genes (2754 in total, **Supplementary File S2**) for 13 previously published citrus genomes and 6 citrus-related genomes, together with HY1 and HY2 were identified and used for phylogeny development. Five calibration points (annotated in tree in **Figure 3A**: 3.49∼7.39, 3.92∼11.0, 12.8∼13.0, 7.57∼25.92, and 13.5∼30.0 Mya) were applied to estimate the divergence time. As shown in **Figure 3A**, the 6 citrus-related species were resolved as basal branches (group F), followed by the early diverging citrus species *P. trifoliata* (group G). The rest citrus species could be divided into 5 major groups (A-E highlighted in **Figure 3A**). Three of those corresponded to the well-recognized citrus ancestors: mandarin (A: *C. reticulata*), pumelo (C: *C. maxima*), and citron (E: *C. medica*). Consistent with the genome tracing results, HY1 and HY2 were grouped with mandarin and pumelo, respectively (**Figure 3A**). The divergence time of HY1 was estimated at 2.0 Mya, earlier than the split between *C. clementina* and *C. reticulata* but after the divergence of *C. sinensis*, while HY2 was estimated to diverge at 2.18 Mya, before the split between *C. grandis* and *C. maxima* and after the branching of *C. hongheensis* (**Figure 3A**). Therefore, HY1 and HY2 represented 2 novel genomes that has not been reported before. Using the earliest diverging *Poncirus trifoliata* as reference, ks peak fitting showed that HY1, HY2, and other Citrus species displayed a single ks peaks at similar position (**Figure 3B**), implying no additional large scale whole genome duplication events. Gene family analyses based on the developed phylogeny revealed more than 3-fold expansion (+345) than contraction (-102) in the branch leading to HY1. In contrast, slightly more expansion (+255) than contraction (-162) was observed in the branch leading to HY2 (**Figure 3A**). The percentage of single copy genes in HY1 and HY2 (6.5%) were relatively lower than that in other citrus species (8.1% ∼ 12.1%), probably caused by genome annotation bias (bar plot in **Figure 3A**). Relatively higher percentage of HY2-specific genes (0.6% or 261) was identified than that in HY1 (0.4% or 158). The numbers of genes for 5 important metabolic pathways (map00900: terpenoid biosynthesis, map00906: carotenoid biosynthesis, map00940: phenylpropanoid biosynthesis, map00941: flavonoid biosynthesis, and map04626: plant-pathogen interaction) related to fruit quality were counted (heatmap in **Figure 3A; Supplementary File S3**). Overall, we observed a general expansion of gene members in all target pathways for citrus species, compared to the basal citrus-related species and the early diverging species *P. trifoliata*, implying a potential human selection impact. Particularly, we noticed that group A and C encompassing HY1 and HY2 contained the most expanded gene members compared to other citrus species, except for *C. maxima* Huazhouyou instead displayed relatively contracted gene families in these pathways. Further analysis is needed to investigate if this is caused by genome annotation bias or not. Noteworthy, a total of 158 and 261 species-specific genes were identified for HY1 and HY2, respectively (**Supplementary File S4**). Enrichment analyses showed that gene ontology (GO) terms related to photosynthesis, signal transduction, and response to stimulus were significantly enriched in the HY1-specific genes (**Figure 3C**). In contrast, no significantly enriched GO term could be detected in the HY2-specific genes.

**Figure 3.**
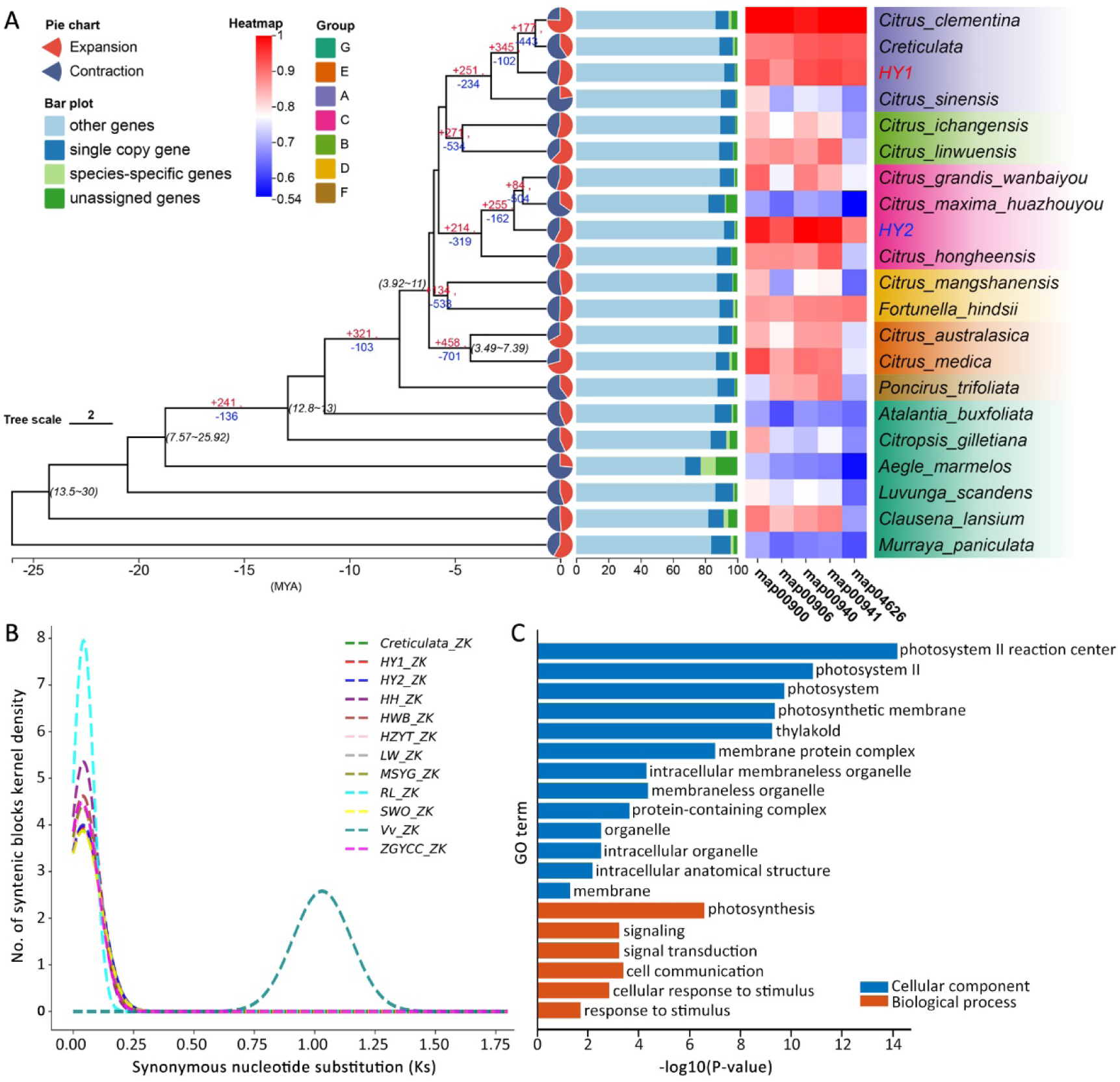
Phylogeny dating and gene family evolution analyses. **A.** Displays the species phylogeny of HY1 and HY2 in references to other citrus species and citrus-related species. The tree topology was obtained using Orthofinder, calibrated at 5 nodes (age annotated in tree) using mcmctree program based on single copy genes. Identified branching groups were highlighted correspondingly. Gene family expansion and contraction at key nodes (numbers) and each taxa (pie charts) were indicated. The percentage of single copy genes, species-specific genes, unassigned genes, and other genes were annotated as bar charts for each taxa. The heatmap represents gene counts for 5 key KEGG pathways: map00900-terpenoid biosynthesis, map00906-carotenoid biosynthesis, map00940-phenylpropanoid biosynthesis, map00941-flavonoid biosynthesis, and map04626-plant-pathogen interaction (Supplementary File S2, S3). **B**. Ks distribution of collinear gene pairs of citrus species with the early diverging *P. trifoliata* (ZK) species as reference (HWB *C. grandis* wanbaiyou, HZYT *C. maxima* huazhouyou, SWO *C. sinensis*, MSYG *C. mangshanensis*, ZGYCC *C. ichangensis*, LW *C. linwuensis*, HH *C. hongheensis*, RL *C. medica*, Vv *Vitis vinifera*). Vitis vinifera was used an outgroup reference. **C**. Gene ontology enrichment analysis of HY1-specific genes. GO terms with corrected p-value <0.05 were included as significant. No significant enrichment was identified for HY2-specific genes (Supplementary File S4).

### Synteny and structural variations analysis of HY1 and HY2

The synteny and structural variations of HY1 and HY2 were inferred in reference to their ancestral genomes: mandarin and pummelo, respectively (**Figure 4A**). Overall, we observed much less large structural variations between mandarin and HY1, compared to that between HY1 and HY2, consistent with our ancestor tracing results that HY1 was derived from mandarin. For example, the large inversions identified between HY1 and HY2 on chromosomes chr6, chr7, and chr8 were not observed between mandarin and HY1, providing direct evidence that HY1 may have originated from a mandarin ancestor. Similar patterns could be found for other chromosomes and other structural variations as well. In contrast to the strong synteny conservation between HY1 and its ancestor genome mandarin, we observed significant large structural variations between HY2 and the pummelo genome (*C. grandis* L. Osbeck.cv. Wanbaiyou) (**Figure 4A**). Noteworthy, these large structural variations such as the inversions on chromosomes chr1, chr5, and chr9 were specific to the pummelo genome used in this study but not present between HY1 and HY2 (**Figure 4A**), implying that these large structural variations between HY2 and pummelo were mainly due to genome re-arrangement in *C. grandis* L. Osbeck.cv. Wanbaiyou but not in the pummelo ancestor. Further analyses are needed to investigate if these large genome re-arrangements are conserved in other pummelo genomes or not.

**Figure 4.**
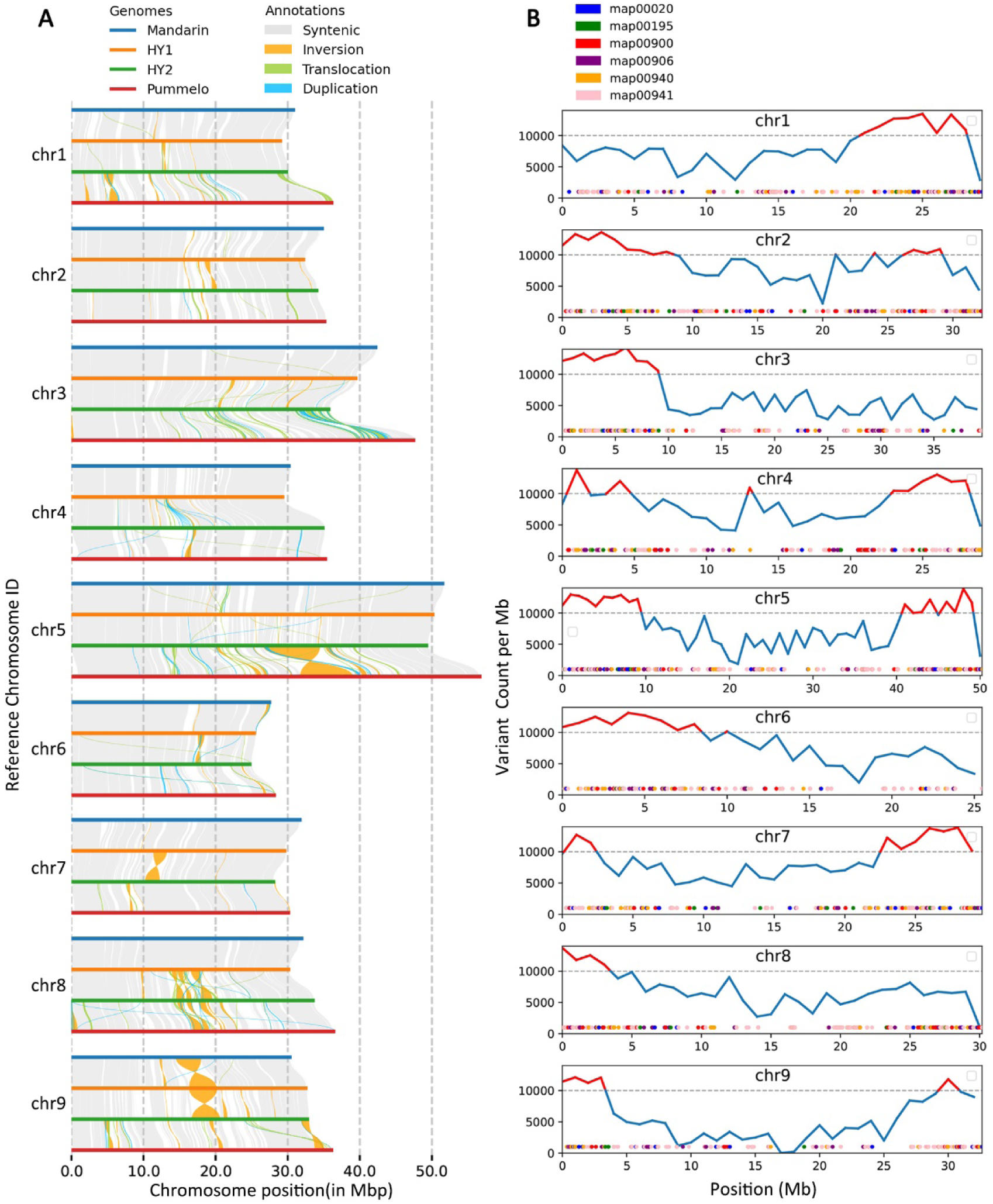
Synteny and genetic variations of HY1, HY2, mandarin, and pummelo genomes. **A.** Display the synteny and structural variations of HY1, HY2, and their ancestral genomes mandarin (*C. reticulata*) and pummelo (*C. grandis*). Pair-wise sequential whole genome alignments in the order of mandarin-HY1-HY2-pummelo were performed. Synteny (grey) and large structural variations (inversions: orange, translocations: green, duplications: blue) were identified using SyRI. **B**. Displays the distribution of the allelic genetic variants (based on SyRI outputs) between HY1 and HY2 and important pathway genes (map00020-citrate cycle, map00195-photosynthesis, map00900-terpenoid biosynthesis, map00906-carotenoid biosynthesis, map00940-phenylpropanoid biosynthesis, map00941-flavonoid biosynthesis).

The genomic variations between HY1 and HY2 were examined specifically to investigate the allelic diversity in Huyou. We identified 1,996,631 SNPs, 150,487 insertions (1.66 Mb), and 139,519 deletions (1.57 Mb) (**Supplementary File S5**). These variations led to the classification of 25,503 highly diverged regions, accounting for (∼ 163.49 Mb or 50%) of the HY1 genome, consistent with heterozygosity. Despite these divergent regions, majority of the genome (252.08 Mb or 77.2%) were found conserved in syntenic regions, implying a generally conserved synteny between the ancestral genomes for HY1 and HY2 before their hybridization that resulted in Huyou. For structural variations, we identified a total of 73 inversions (21.04 Mb), 592 translocations (6.35 Mb), and 938 duplications (8.14 Mb), which were mainly located in the central region of each chromosome (**Supplementary File S5**). Three large inversions between HY1 and HY2 were observed in the middle regions of chr9 and chr8, followed by a few moderate inversions on chr1, chr2, chr3, chr4 and chr6. In addition, a genomic region in the middle of chr5 in HY2 seems to be expanded through significant duplications, which may have contributed to the chromosome extension of chr5 in HY2 compared to chr5 in HY1 (**Figure 4A**). Lastly, the number of genetic variations between HY1 and HY2 were calculated per 1 Mb window along each chromosome. The obtained variation distribution profile revealed a relatively higher-level of genetic diversity in the distal regions of each chromosome compared to the central regions (**Figure 4B**). Mapping the genes from the 6 important metabolic pathways (map00020-Citrate cycle, map00195-Photosynthesis, map00900-terpenoid biosynthesis, map00906-carotenoid biosynthesis, map00940-phenylpropanoid biosynthesis, map00941-flavonoid biosynthesis) to the genome showed that many of these genes were located within the identified divergent regions, suggesting a high allelic diversity for important fruit quality characters in Huyou.

### Allelic imbalance of gene expression profiles in different Huyou tissues

As the most heterozygous citrus genome to date, Huyou provides an invaluable opportunity to study genome evolution. To investigate the allelic imbalance post ancient hybridization, we retrieved public transcriptome data of 6 different tissues (root, stem, leaf, flower, unripe fruit, ripe fruit) in Huyou from a previous study (Miao*, et al.*, 2024) and compared the expression levels of collinear genes (26,092 in total) between HY1 and HY2. The percentages of genes preferentially expressed in HY1 (HY1higher (log_2_^(HY1/HY2)^ ≥ 1) and HY2 (HY2higher log_2_^(HY1/HY2)^ ≤ -1) were calculated across each chromosome and tissue (**Supplementary File S6**). Overall, we observed relatively higher numbers of HY2-preferential genes than HY1-preferential genes across most tissues and chromosomes (**Figure 5A**), implying a clear transcriptional bias for HY2 genes following hybridization. Noteworthy, we found that the HY2 allelic dominance was exceptionally prominent in root tissue (HY2higher at 12.5% versus HY1higher at 7.24%) compared to that in the other tissues (leaf, stem, flower, unripe fruit, ripe fruit), which only displayed slight HY2 allelic dominance (11.67% versus 11.10% on average), implying a strong tissue-specific pattern. The exceptional HY2 allelic dominance in root tissue across the 9 chromosomes could be seen in **Figure 5B**, relatively more evident on chromosomes chr1-7 and relatively mild on chr8 and chr9. This might be caused by the large segmental replacements observed on chr8 and chr9. Indeed, at the chromosome level, chr9 displayed the lowest level of total allelic divergence (HY1higher + HY2higher = 16.77%) and followed by chr8 (21.29%) and other chromosomes (22.30-24.27%).

**Figure 5.**
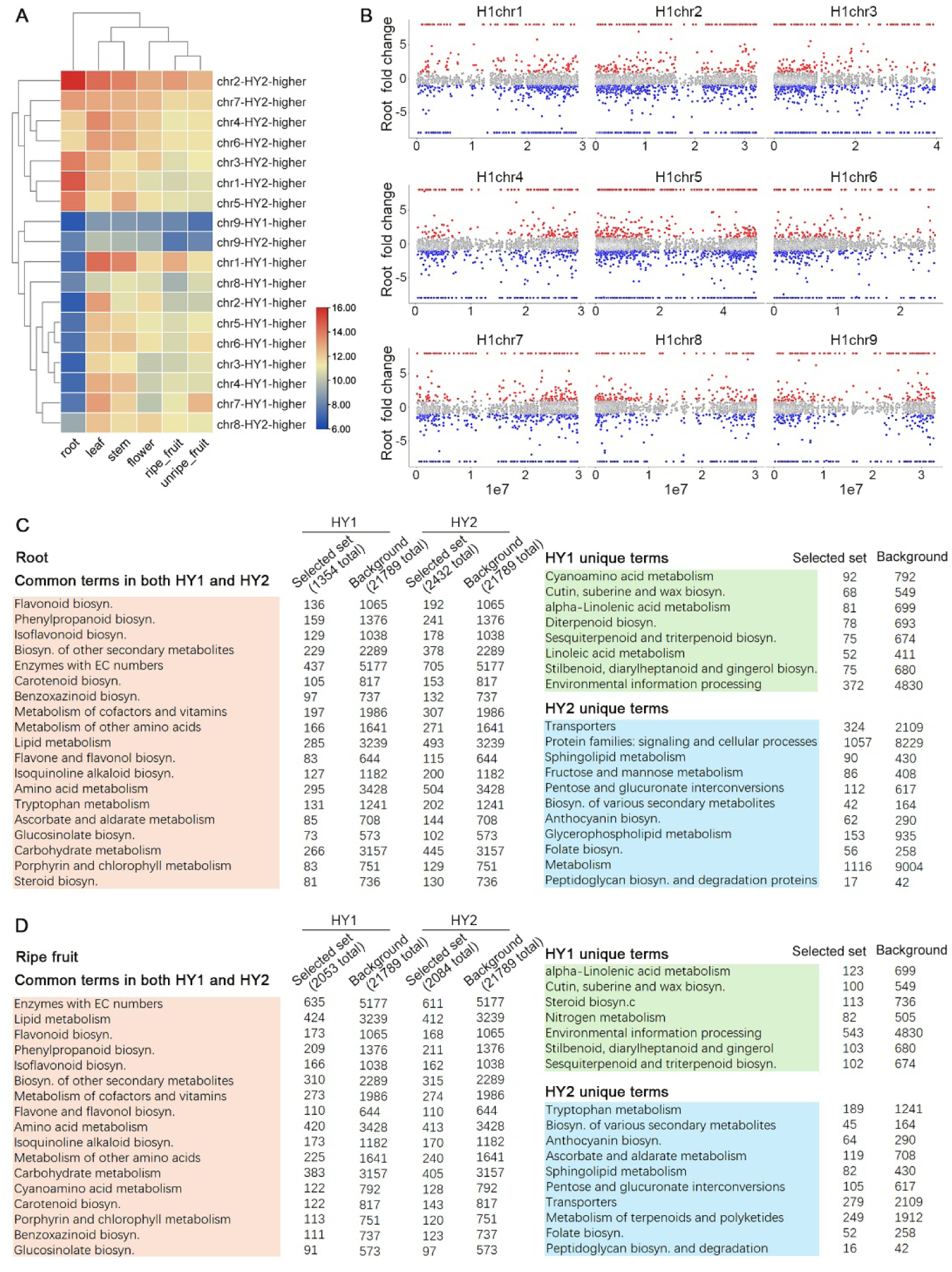
Transcriptional alleic imbalance between HY1 and HY2 in different tissues. **A.** Clustering of the percentages of HY1-preferential (log_2_^(HY1/HY2)^ ≥ 1) and HY2-preferential (log_2_^(HY1/HY2)^ ≤ -1) genes 6 tissues (root, stem, leaf, flower, unripe fruit, ripe fruit) in each chromosome (**Supplementary File S7**). Only the collinear genes between HY1 and HY2 were counted. The percentages of HY1 and HY2 preferential genes were calculated relative to the total number of gene in each chromosome. **B**. Displays the distribution of HY1 and HY2 preferential genes along each chromosome for the root transcriptome. **C**. KEGG enrichment of allelic imbalance collinear genes in root with corrected p-value <1 × 10^-5^ were shown. D. KEGG enrichment of allelic imbalance collinear genes in ripe fruit with corrected p-value <1 × 10^-5^ were shown.

Due to the exceptional HY2 allelic dominance in the root tissue, the collinear genes with transcriptional preference from HY1 (1354 HY1higher genes) and HY2 (2432 HY2higher genes) were used for KEGG pathway enrichment analysis (corrected p-value ≤ 1-e5). Interestingly, these two sets of genes shared some commonly enriched pathways that are mainly related to antioxidants biosynthesis, including flavonoid, phenylpropanoid, isoflavonoid, carotenoid, benzoxazinoid, flavone, flavonol, alkaloid, ascorbate, and steroid biosynthesis (**Figure 5C**). It is well-known that antoxidant biosynthesis plays critical role in plants’ adaptation to various biotic and abiotic stresses. The allelic imbalance of genes within these pathways in the root tissue may have important implication in Huyou’s environmental adaptation following its ancient genome hybridization. In addition to these common pathways, HY1 preferential genes were uniquely enriched with amino acid and linoleic acid metabolism and the biosynthesis of cutin, wax, diterpenoid, triterpenoid, sesquiterpenoid, stilbenoid, diarylheptanoid, and gingerol which are closely linked to environmental adaptation (**Figure 5C**). In contrast, HY2 preferential genes were uniquely enriched with protein families related to signaling and cellular processes, including transporter genes, sphingolipid metabolism, and genes responsible for fructose, mannose, pentose, glucuronate interconversions, glycerolphospholipid, peptidoglycan metabolism. In addition, the biosynthesis of anthocyanin, folate, and various secondary metabolites were also enriched in HY2 preferential genes but not in HY1 preferential genes.

In addition to the root tissue, the allelic imbalance genes from HY1 (2053HY1higher genes) and HY2 (2084 HY2higher genes) in ripe fruit tissue were also investigated. Overall, we observed a strikingly similar enrichment profiles with that in the root tissue (**Figure 5D**), suggesting that the allelic imbalance at the gene functional level may be conserved across tissues despite the variation in gene numbers. In details, like that in the root tissue, the commonly enriched pathways from HY1 and HY2 were also prominently related to antioxidants biosynthesis such as flavonoid, phenylpropanoid, isoflavonoid, flavone, flavonol, isoquinoline alkaloid, carotenoid, benzoxazinoid, and glucosinolate. Other common pathway terms included lipid, cofactors and vitamin, amino acid, porphyrin and chlorophyll metabolism which were also detected in the root tissue (**Figure 5D**). The unique pathway terms in HY1 (linolenic acid metabolism and the biosynthesis of cutin, suberine, wax, stilbenoid, diarylheptanoid, gingerol, sesquiterpenoid, triterpenoid) and HY2 (tryptophan, ascorbate, aldarate, sphingolipid, pentose, glucuronate, terpenoids metabolisms and the biosynthesis of secondary metabolites, anthocyanin, folate, peptidoglycan) in ripe fruit tissue were also similar to that in the root tissue (**Figure 5D**).Tissue-specific allelic imbalance in key pathways

Carotenoid biosynthesis pathway (map00906) plays a critical role in citrus fruit quality, controlling fruit color and flavor. To investigate the genomic characteristics of key Huyou candidate genes in this pathway, we first investigated the allelic imbalance by mapping the ratio of HY1 and HY2 collinear gene transcription to this pathway. Whilst the major components of the pathway displayed no allelic preference, we observed a clear tissue-specific allelic imbalance for some key genes responsible for the astaxanthin biosynthesis specifically (**Figure 6**). For instance, we found one enzyme (EC 1.14.15.24) encoding beta-carotene 3-hydroxylase displayed a consistent HY2 dominance in the root tissue but a HY1 dominance in the stem, flower, unripe fruit, and ripe fruit tissue and no imbalance in leaf. Another enzyme (EC 1.23.5.1) encoding violaxanthin de-epoxidase displayed HY1 dominance in root, leaf, ripe fruit but no imbalance in the other tissue. Conversely, the enzyme (EC 1.2.3.14) encoding abscisic-aldehyde oxidase display HY2 dominance in stem, leaf, and unripe fruit but not in other tissue. These observations provided direct support of various allelic imbalance present in Huyou genome and shed valuable insights into genome evolution following ancient hybridization.

**Figure 6.**
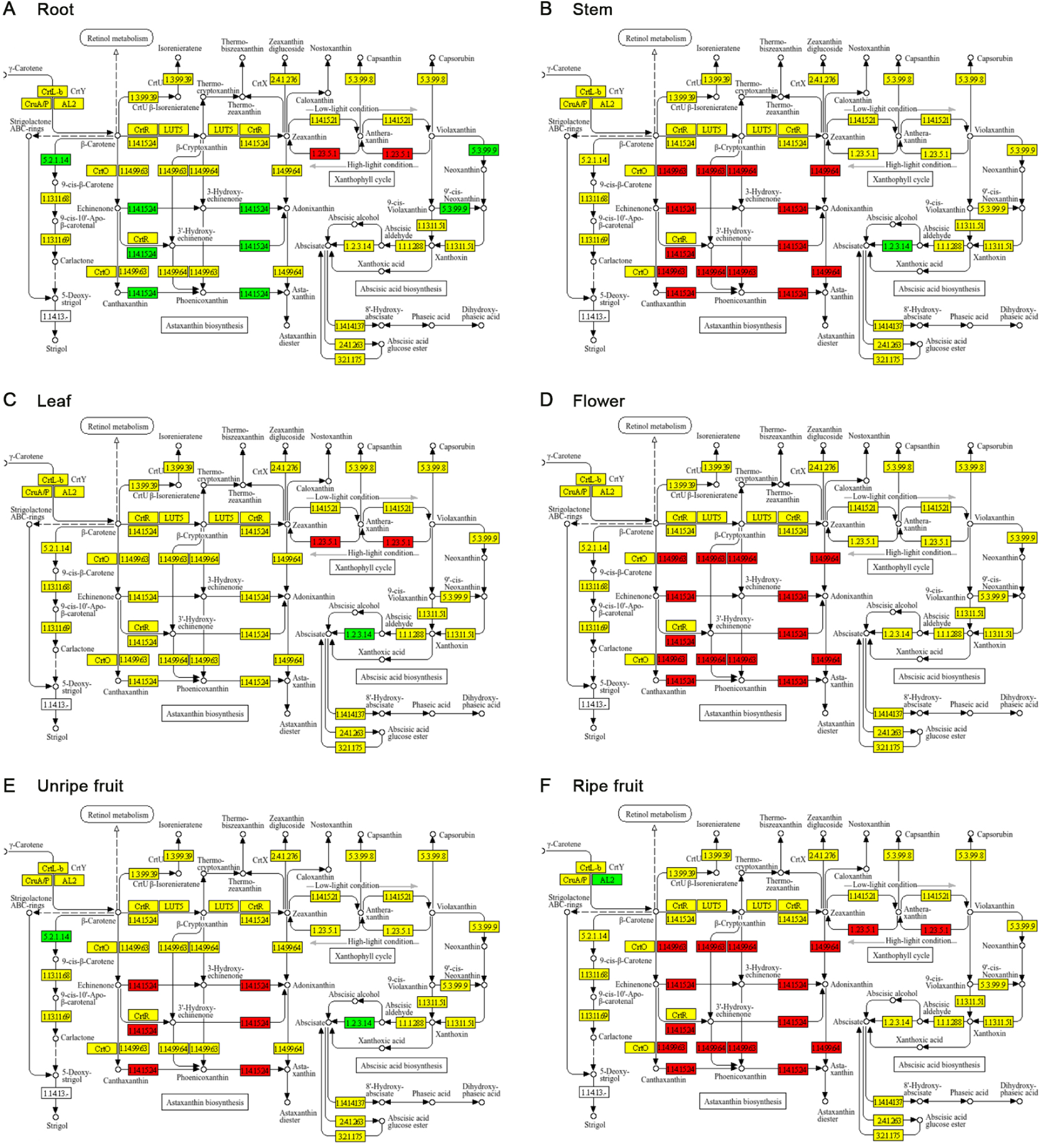
Displays the transcriptome allelic imbalance of HY1 and HY2 for map00906 pathway. The ratios of HY1 and HY2 collinear genes (log2^(HY1/HY2)^) were mapped to KEGG pathway map00906 using TBtools. Transcriptome data in 6 Huyou tissues (root, stem, leaf, flower, unripe fruit and ripe fruit) from a previous study was downloaded and analyzed. Yellow, red, and green highlights indicated neutral, HY1 dominant, and HY2 dominant, respectively. Only the Astaxanthin biosynthesis module of map00906 were displayed.

To characterize the complete profile of the carotenoid biosynthesis pathway in Huyou, we further extracted the gene copy number information of key genes (represented by 39 KEGG orthologs) in HY1, HY2, and another 8 citrus species and explored their transcriptional profiles in 6 Huyou tissues. Overall, we observed conserved gene presence and copy number pattern across different citrus species, with cytochrome P450 (CYP701, CYP97C1, CYP707A, CYP97A3, CYP82G1) as the most expanded protein families within the carotenoid pathway. Copy number variations (CNVs) were observed in all KO orthologs except for 2 single copy genes K15744 (zeta-carotene isomerase) and K17911 (beta-carotene isomerase) (**Figure 7A**). Similarly, HY1 and HY2 displayed varied copy numbers in 33 out of the 39 KO orthologs. Noteworthy, Huyou (HY1 or HY2) displayed the highest copy numbers in 21 out of the 39 KO orthologs (**Figure 7A**). At the gene expression level, we observed varied transcriptional profiles across the 6 tissues investigated (**Figure 7B**). Interestingly, we found that over half of the KO orthologous genes in Huyou displayed the highest expression in the flower tissue, followed by the root tissue. Instead, out of the 39 KO orthologs, we found 17 KOs displayed above median transcription in fruit tissues (either unripe or ripe), of which 6 KOs (FDFT1, LCYB, LCYE, CCS1, AOG, DWARF27) displayed the highest expression in fruit tissues (highlighted in red rectangles in **Figure 7C**), which require more attention for their potential critical roles in fruit development, despite that some of these genes may not be expanded in Huyou (**Figure 7C**). At the pathway map level, we found most of the genes leading to the terpenoid biosynthesis (C30, C15, C5, C10, C20) were found expanded compared to other species, whilst this pattern was not prominent for the carotenoid biosynthesis (C40) (**Figure 7C**). Taken together, CNV and transcriptome analyses revealed unique genomic characteristics for the carotenoid biosynthesis pathway in Huyou which may have contributed to its distinct fruit flavor against other citrus species.

**Figure 7.**
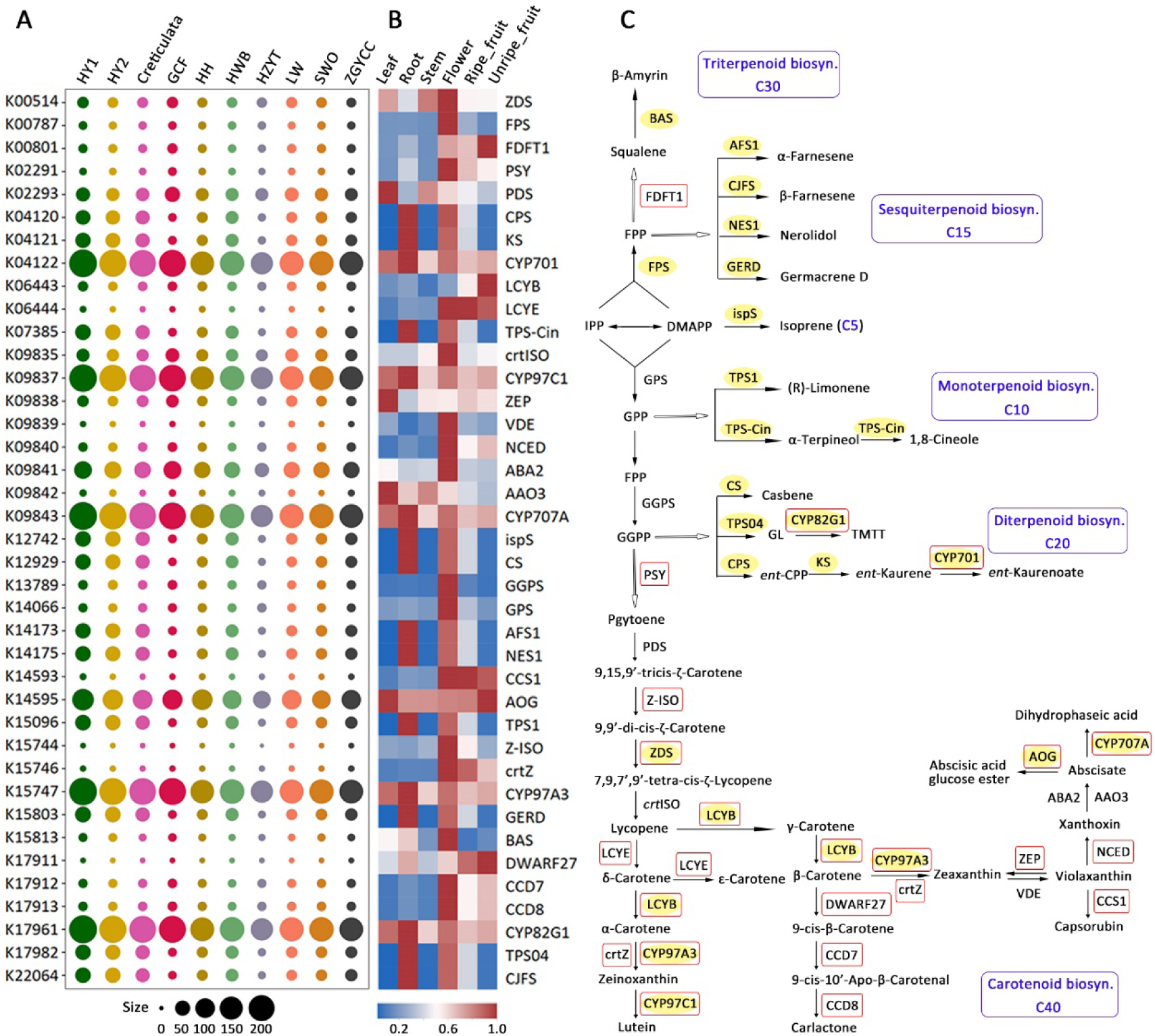
The CNVs and transcriptome profiles for carotenoid pathway in Huyou. **A.** Displays the CNV in HY1, HY2, and other 8 citrus species included in this study. Gene counts were based on results from Orthofinder analysis in Figure 3. Genes were assigned to KO orthologs based on kegg annotation. **B**. Heatmap displaying the transcription level of each KO ortholog in 6 Huyou tissues. The transcriptome values for multiple genes were summed and normalized (divided by maximum value) across the 6 tissues. **C**. Display the metabolic map of the carotenoid pathway. Genes whose copy numbers are highest in Huyou (either HY1 or HY2) were highlighted in yellow. Genes with above median expression in fruit tissues were enclosed in red rectangles.

## Discussion

As a unique natural hybrid citrus landrace first discovered and widely grown in Zhejiang province in China, Huyou holds important agricultural, economic, and medicinal values due to its premium fruit quality characteristics such as a golden skin, abundant bioactive components, extended storage life. In this study, we leverage PacBio long-read and Hi-C sequencing technologies and presented 2 high-quality haplotype-resolved genome assemblies HY1 and HY2 of 120 years old Huyou “ancestral tree”. Despite the recent availability of another Huyou haplotype-resolved genome data (Miao*, et al.*, 2024), our in-depth ancestor tracing, phylogeny dating, genome structure, and transcriptome analyses offered an unprecedent view of Huyou’s genetic makeup and evolutionary origin. The explicit determination of Huyou’s genome origin from a mandarin and pummelo hybridization event solved a long biological mystery that has puzzled local Huyou growers and botanists (Li*, et al.*, 2019; Chen*, et al.*, 2002; Chen*, et al.*, 2006; Xu*, et al.*, 2006). Based on whole genome survey of 58 worldwide diverse citrus germplasms, hybrid citrus species derived from 2 or more ancestral species have been found to have generally high genome heterozygosity at 1.5-2.4% (Wu*, et al.*, 2018). Here, we highlight the exceptional genome heterozygosity of Huyou at 3.07%, among the highest across known citrus genomes, which will make Huyou an interesting subject for genome evolution studies by broad citrus researchers in the future. The haplotype genomes HY1 and HY2 obtained in this study essentially represent 2 novel ancestral citrus genomes not reported before, thus would significantly contribute to the currently limited citrus genetic pool. This study will enable citrus breeders to harness the genetic potential of Huyou more effectively, potentially accelerating the development of improved citrus varieties with unique traits.

The explicit resolution of Huyou’s genome origin in this study highlights the power of haplotype-resolved genomes in dissecting the complex evolutionary origin of citrus species. In the citrus genus, there are a total of 30 species, most of which have been derived from the hybridization across 3 ancestral species: mandarin (*C. reticulata*), pummelo (*C. maxima*), citron (*C. medica*). The ancient hybrids were subjected to further reticulate hybridization during citrus domestication, leading to a complex history of admixture in current citrus cultivars and landraces characterized by high heterozygosity (Wu*, et al.*, 2018). Tracing the evolution and domestication history of citrus species has been extremely challenging in traditional citrus genome studies based on short read sequencing, which often produced a mosaic genome assembly and failed to account for allelic variants, leading to incomplete or inaccurate representations of the genome. By leveraging a k-mer-based tracing approach on the haplotype-resolved genomes, we are able to resolve and quantify the ancestral components of Huyou to specific genomic regions. In addition, the haplotype genome tracing also allows us to dissect the segmental insertions, deletions, inversions, recombinations and other structural variations between HY1 and HY2, particularly the recombination events on chr8 and chr9 in HY2. These unprecedented genomic insights are critical for the accurate interpretation of the genomic and phenotypic features of Huyou. Our study was inspired by a recent study in which the haplotype-resolved lemon genome was traced to be an admixture of 3 ancestor genomes of citron, mandarin, and pummelo (Bao*, et al.*, 2023). Notably, numerous studies highlight the increasing trend of generating chromosome-level and haplotype-resolved genome assemblies across the plant kingdom, such as potato, lychee, tea, and strawberry. The advantages of haplotype-resolved genome for citrus genome are being increasingly appreciated in several recent citrus genome research including *C. limon* (Mario*, et al.*, 2021), *C. australis* (Nakandala*, et al.*, 2023), *C. changshanensis* (Miao*, et al.*, 2024), *Papeda* (Wang*, et al.*, 2025). However, we noticed that only the collapsed genome sequence was reported and analyzed in several haplotype-resolved genome studies, which in our opinion should be avoided, particularly for those organisms with high heterozygosity such as Citrus species. We expect an increasingly more haplotype-resolved citrus genomes to be generated and published, which is positioned to revolutionize the citrus genome evolution and domestication research.

Allelic imbalance (AI) refers to the unequal expression of alleles from a heterozygous locus, leading to differential gene expression. This is another critical insight gained from haplotype-resolved genome assembly. In this study, based on gene expression analysis of collinear genes between HY1 and HY2, we observed clear evidence of AI (HY2 dominant over HY1) across all 6 tissues investigated, suggesting AI may be a prevalent phenomenon in genome evolution in the plant kingdom. One of the most notable findings in this study is that we observed AI tends to be imbalanced across tissues, i.e the HY2 allelic dominance was much more prominent in some tissues such as root over the others. Metabolic pathway enrichment analyses of AI genes in Huyou root tissue revealed a strong over-representation of genes related to various antioxidant production, such as flavonoid, phenylpropanoid, isoflavonoid, carotenoid, benzoxazinoid, flavone, flavonol, alkaloid, ascorbate, and steroid biosynthesis. These antioxidants have well-known biological functions in plants’ adaptation to various biotic and abiotic stresses. The allelic imbalance of genes within these pathways in the root tissue may have important implication in Huyou’s environmental adaptation following its ancient genome hybridization. It is interesting to observe that these antioxidant biosynthesis terms were over-represented in both HY1-higher and HY2-higher genes, suggesting an overall evolutionary preference of AI genes related to environmental adaptation. At the plant tissue level, we observed overall conservation of AI functions across tissues such as root and ripe fruit, despite a significantly higher degree of AI in root. This novel insight needs to be verified in future analysis in other species. In plants, especially those with high heterozygosity, AI plays a significant role in genome evolution, influencing traits such as disease resistance, growth patterns, and adaptation to environmental changes (Jang*, et al.*, 2024; Djari*, et al.*, 2024; Li*, et al.*, 2023). The impact of AI on plants’ genome evolution and environmental adaption has been well recognized in several recent haplotype-resolved genome studies. For example, a study in *Populus tomentosa* identified heterozygous alleles associated with adaptive traits, highlighting the role of AI in evolutionary processes (Li*, et al.*, 2023). Similarly, haplotype-resolved assemblies grapevine identified structural variations between haplotypes associated with allelic expression linked to disease resistance and fruit quality (Djari*, et al.*, 2024). Another recent study in *Pinus densiflora* revealed the functional contributions of AI to flowering and abiotic stress-related traits (Jang*, et al.*, 2024). In addition to the overlapping AI functions, haplotype-specific AI functions were also observed. The biological implications of our observed AI patterns remain to be characterized, particularly for citrus genome research. As far as the authors are concerns, our study is the first to demonstrate the presence of strong AI in citrus hybrid genome. Based on our observation of AI in Huyou, we reason that AI should be prevalent across the various citrus genomes due to the presence of frequent hybridization events. Given the still limited haplotype resolved genome assemblies for citrus species, the importance of AI in citrus genome evolution and its interplay with phenotypic diversity in citrus deserves much more attention in future research.

## Conclusions

We sequenced a 120-year-old "ancestral tree" of *C. changshan* Y. B. Chang (Huyou), a unique natural hybrid citrus landrace first discovered in Changshan, Zhejiang province, China. Huyou displayed genome heterozygosity at 3.07%, among the highest across known citrus genomes. We presented two high-quality haplotype-resolved genomes HY1 and HY2 and determined that Huyou had originated from a mandarin and pummelo hybridisation event, resolving a mystery that has puzzled local Huyou growers and botanists. HY1 and HY2 had diverged at 2.0 Mya and 2.18 Mya, earlier than the split of *C. clementina* and *C. reticulata*, and the split of *C. grandis* and *C. maxima*, respectively. Haplotype-based transcriptome analyses also demonstrated the presence of strong tissue-specific allelic imbalance in Huyou’s genome. The haplotype genomes HY1 and HY2 represent two novel ancestral citrus genomes not previously reported, contributing to the limited citrus genetic pool and enabling citrus breeders to harness Huyou’s genetic potential. The explicit resolution of Huyou’s genome origin highlights the power of haplotype-resolved genomes in dissecting the complex evolutionary origin of citrus species.

## Competing interests

The authors declare no competing interest.

## Acknowledgements

We acknowledge the relevant citrus research communication for making the citrus genomic data available to the public, particularly the recently published Huyou transcriptome data (Miao*, et al.*, 2024).

## Author contributions

YJ, CL, ZZ designed and supervised the study. ZZ, YJ, TX, YL, HH, LL, BG, PZ, CT, TQ, ZY, ZC performed data analysis. YJ, ZZ, XH, YL wrote the manuscript. TX, CL, LW edited the manuscript. All authors have read and approved the manuscript.

## Data availability

The raw sequencing data has been deposited at NCBI with project ID PRJNA1090545 (release upon acceptance). The genome assemblies and gene annotations are available at https://figshare.com/s/ba2b2081a2512770c31a

## Funding

This work was supported by the National Natural Science Foundation of China (32301872), the Natural Science Foundation of Zhejiang Province (LQ23C130003), and Interdisciplinary Research Project of Hangzhou Normal University (2025JCXK01).

## Supplementary Information

Supplementary File S0. TE annotation for HY1 and HY2.

Supplementary File S1. k-mer tracing results of HY1 and HY2.

Supplementary File S2. Orthofinder gene matrix for target species in this study.

Supplementary File S3. List of Orthogroup IDs identified for key KEGG pathways.

Supplementary File S4. List of species-specific gene in HY1 and HY2.

Supplementary File S5. SyRI summary for HY1 and HY2 synteny analysis.

Supplementary File S6. RNAseq results (quantified TPM and HY1/HY2 ratio) HY1 and HY2 collinear genes in six tissues.

Supplementary File S7. Percentage of allelic imbalance genes in each chromosome across 6 tissues.

## Notes

### Competing Interest Statement

The authors have declared no competing interest.

## References

Gmitter, FG, Chen, C, Machado, MA, de Souza, AA, Ollitrault, P, Froehlicher, Y, Shimizu, T. 2012. Citrus genomics. Tree Genetics & Genomes, 8: 611–626.

Wu, GA, Terol, J, Ibanez, V, López-García, A, Pérez-Román, E, Borredá, C, Domingo, C, Tadeo, FR, Carbonell-Caballero, J, Alonso, R et al. 2018. Genomics of the origin and evolution of Citrus. Nature, 554: 311.

Li, Y, Zhou, S, Tong, X, Wang, L, Xiang, T. 2019. Analysis of the Origin of *Citrus changshan-huyou* by DNA Barcode. International Journal of Agriculture and Biology, 22: 973–978.

Chang, Y-b. 1991. A new species of genus citrus from China. Bulletin of Botanical Research, 11: 5–7.

Zhang, J, Sun, C, Yan, Y, Chen, Q, Luo, F, Zhu, X, Li, X, Chen, K. 2012. Purification of naringin and neohesperidin from Huyou (*Citrus changshanensis*) fruit and their effects on glucose consumption in human HepG2 cells. Food chemistry, 135: 1471–1478.

Guo, J-j, Gao, Z-p, Xia, J-l, Ritenour, MA, Li, G-y, Shan, Y. 2018. Comparative analysis of chemical composition, antimicrobial and antioxidant activity of citrus essential oils from the main cultivated varieties in China. Lwt-Food Science and Technology, 97: 825–839.

Sheng, L, Shen, D, Luo, Y, Sun, X, Wang, J, Luo, T, Zeng, Y, Xu, J, Deng, X, Cheng, Y. 2017. Exogenous gamma-aminobutyric acid treatment affects citrate and amino acid accumulation to improve fruit quality and storage performance of postharvest citrus fruit. Food chemistry, 216: 138–145.

Jiang, J, Yan, L, Shi, Z, Wang, L, Shan, L, Efferth, T. 2019. Hepatoprotective and anti-inflammatory effects of total flavonoids of *Qu Zhi Ke* (peel of *Citrus changshan-huyou*) on non-alcoholic fatty liver disease in rats via modulation of NF-kappaB and MAPKs. Phytomedicine, 64: 153082.

Miao, C, Wu, Y, Wang, L, Zhao, S, Grierson, D, Xu, C, Chen, W, Chen, K. 2024. Haplotype-resolved chromosome-level genome assembly of Huyou (*Citrus changshanensis*). Scientific Data, 11: 605.

Xu, Q, Chen, LL, Ruan, XA, Chen, DJ, Zhu, AD, Chen, CL, Bertrand, D, Jiao, WB, Hao, BH, Lyon, MP et al. 2013. The draft genome of sweet orange (*Citrus sinensis*). Nature Genetics, 45: 59–68.

Wu, GA, Prochnik, S, Jenkins, J, Salse, J, Hellsten, U, Murat, F, Perrier, X, Ruiz, M, Scalabrin, S, Terol, J et al. 2014. Sequencing of diverse mandarin, pummelo and orange genomes reveals complex history of admixture during citrus domestication. Nature Biotechnology, 32: 656–663.

Wang, X, Xu, YT, Zhang, SQ, Cao, L, Huang, Y, Cheng, JF, Wu, GZ, Tian, SL, Chen, CL, Liu, Y et al. 2017. Genomic analyses of primitive, wild and cultivated citrus provide insights into asexual reproduction. Nature Genetics, 49: 765–774.

Bao, Y, Zeng, Z, Yao, W, Chen, X, Jiang, M, Sehrish, A, Wu, B, Powell, CA, Chen, B, Xu, J et al. 2023. A gap-free and haplotype-resolved lemon genome provides insights into flavor synthesis and huanglongbing (HLB) tolerance. Hortic Research, 10: uhad020.

Peng, Z, Bredeson, JV, Wu, GHA, Shu, SQ, Rawat, N, Du, DL, Parajuli, S, Yu, QB, You, Q, Rokhsar, DS et al. 2020. A chromosome-scale reference genome of trifoliate orange (*Poncirus trifoliata*) provides insights into disease resistance, cold tolerance and genome evolution in. Plant Journal, 104: 1215–1232.

Huang, Y, He, JX, Xu, YT, Zheng, WK, Wang, SH, Chen, P, Zeng, B, Yang, SZ, Jiang, XL, Liu, ZS et al. 2023. Pangenome analysis provides insight into the evolution of the orange subfamily and a key gene for citric acid accumulation in citrus fruits. Nature Genetics, 55: 1964–1975.

Liu, HMZ, Wang, X, Liu, SJ, Huang, Y, Guo, YX, Xie, WZ, Liu, H, ul Qamar, MT, Xu, Q, Chen, LL. 2022. Citrus Pan-Genome to Breeding Database (CPBD): A comprehensive genome database for citrus breeding. Molecular Plant, 15: 1503–1505.

Humann, J, Crabb, J, Frank, M, Cheng, CH, Zheng, P, Lee, T, Buble, K, Hough, H, Scott, K, Jung, S et al. 2022. Using the Citrus Genome Database for Genetics, Genomics, and Breeding Research. Hortscience, 57: S95.

Mario, D, Marco, M, Mirko, M, Chiara, C, Michela, T, Deng, ZN, Alessandro, C, Marco, C, Gaetano, D, Stefano, L et al. 2021. The haplotype-resolved reference genome of lemon (*Citrus limon* L. Burm f.). Tree Genetics & Genomes, 17: 46.

Nakandala, U, Masouleh, AK, Smith, MW, Furtado, A, Mason, P, Constantin, L, Henry, RJ. 2023. Haplotype resolved chromosome level genome assembly of Citrus australis reveals disease resistance and other citrus specific genes. Horticulture Research, 10.

Wang, F, Wang, S, Wu, Y, Jiang, D, Yi, Q, Zhang, M, Yu, H, Yuan, X, Li, M, Li, G et al. 2025. Haplotype-resolved genome of a papeda provides insights into the geographical origin and evolution of Citrus. Journal of integrative plant biology, 67: 276–293.

Zhang, H, Wang, F, Zeng, C, Zhu, W, Xu, L, Wang, Y, Zeng, J, Fan, X, Sha, L, Wu, D et al. 2022. Development and application of specific FISH probes for karyotyping Psathyrostachys huashanica chromosomes. BMC Genomics, 23: 309.

Li, Y, Zhou, S, Feng, M, Lin, Y, Li, L, Ma, G, Xiang, T. 2017. Regeneration of plants from ancestry tree of *Citrus changshan-huyou* Y. B. Chang via tissue culture. Bangladesh Journal of Botany, 46: 1233–1240.

Ranallo-Benavidez, TR, Jaron, KS, Schatz, MC. 2020. GenomeScope 2.0 and Smudgeplot for reference-free profiling of polyploid genomes. Nature Communications, 11: 1432.

Strijk, JS, Hinsinger, DD, Roeder, MM, Chatrou, LW, Couvreur, TLP, Erkens, RHJ, Sauquet, H, Pirie, MD, Thomas, DC, Cao, K. 2021. Chromosome-level reference genome of the soursop (*Annona muricata*): A new resource for Magnoliid research and tropical pomology. Molecular Ecology Resources, 21: 1608–1619.

Chen, S, Zhou, Y, Chen, Y, Gu, J. 2018. fastp: an ultra-fast all-in-one FASTQ preprocessor. Bioinformatics, 34: i884–i890.

Cheng, HY, Concepcion, GT, Feng, XW, Zhang, HW, Li, H. 2021. Haplotype-resolved de novo assembly using phased assembly graphs with hifiasm. Nature Methods, 18: 170–177.

Guan, D, McCarthy, SA, Wood, J, Howe, K, Wang, Y, Durbin, R. 2020. Identifying and removing haplotypic duplication in primary genome assemblies. Bioinformatics, 36: 2896–2898.

Li, H. 2018. Minimap2: pairwise alignment for nucleotide sequences. Bioinformatics, 34: 3094–3100.

Zhang, H, Song, L, Wang, X, Cheng, H, Wang, C, Meyer, CA, Liu, T, Tang, M, Aluru, S, Yue, F et al. 2021. Fast alignment and preprocessing of chromatin profiles with Chromap. Nature Communications, 12: 6566.

Zhou, C, McCarthy, SA, Durbin, R. 2023. YaHS: yet another Hi-C scaffolding tool. Bioinformatics, 39: btac808.

Durand, NC, Robinson, JT, Shamim, MS, Machol, I, Mesirov, JP, Lander, ES, Aiden, EL. 2016. Juicebox Provides a Visualization System for Hi-C Contact Maps with Unlimited Zoom. Cell Systems, 3: 99–101.

Alonge, M, Lebeigle, L, Kirsche, M, Jenike, K, Ou, S, Aganezov, S, Wang, X, Lippman, ZB, Schatz, MC, Soyk, S. 2022. Automated assembly scaffolding using RagTag elevates a new tomato system for high-throughput genome editing. Genome Biology, 23: 258.

Simao, FA, Waterhouse, RM, Ioannidis, P, Kriventseva, EV, Zdobnov, EM. 2015. BUSCO: assessing genome assembly and annotation completeness with single-copy orthologs. Bioinformatics, 31: 3210–3212.

Baril, T, Galbraith, J, Hayward, A. 2024. Earl Grey: A Fully Automated User-Friendly Transposable Element Annotation and Analysis Pipeline. Molecular Biology and Evolution, 41: msae068.

Keilwagen, J, Hartung, F, Grau, J. 2019. GeMoMa: Homology-Based Gene Prediction Utilizing Intron Position Conservation and RNA-seq Data. Methods in Molecular Biology, 1962: 161–177.

Dobin, A, Davis, CA, Schlesinger, F, Drenkow, J, Zaleski, C, Jha, S, Batut, P, Chaisson, M, Gingeras, TR. 2013. STAR: ultrafast universal RNA-seq aligner. Bioinformatics, 29: 15–21.

Chan, PP, Lin, BY, Mak, AJ, Lowe, TM. 2021. tRNAscan-SE 2.0: improved detection and functional classification of transfer RNA genes. Nucleic Acids Research, 49: 9077–9096.

Nawrocki, EP, Kolbe, DL, Eddy, SR. 2009. Infernal 1.0: inference of RNA alignments. Bioinformatics, 25: 1335–1337.

Chen, C, Chen, H, Zhang, Y, Thomas, HR, Frank, MH, He, Y, Xia, R. 2020. TBtools: An Integrative Toolkit Developed for Interactive Analyses of Big Biological Data. Molecular Plant, 13: 1194–1202.

Krzywinski, M, Schein, J, Birol, I, Connors, J, Gascoyne, R, Horsman, D, Jones, SJ, Marra, MA. 2009. Circos: an information aesthetic for comparative genomics. Genome Research, 19: 1639–1645.

Wang, L, He, F, Huang, Y, He, J, Yang, S, Zeng, J, Deng, C, Jiang, X, Fang, Y, Wen, S et al. 2018. Genome of Wild Mandarin and Domestication History of Mandarin. Molecular Plant, 11: 1024–1037.

Zheng, W, Zhang, W, Liu, D, Yin, M, Wang, X, Wang, S, Shen, S, Liu, S, Huang, Y, Li, X et al. 2023. Evolution-guided multiomics provide insights into the strengthening of bioactive flavone biosynthesis in medicinal pummelo. Plant Biotechnology Journal, 21: 1577–1589.

Goel, M, Sun, H, Jiao, WB, Schneeberger, K. 2019. SyRI: finding genomic rearrangements and local sequence differences from whole-genome assemblies. Genome Biology, 20: 277.

Jia, KH, Wang, ZX, Wang, L, Li, GY, Zhang, W, Wang, XL, Xu, FJ, Jiao, SQ, Zhou, SS, Liu, H et al. 2022. SubPhaser: a robust allopolyploid subgenome phasing method based on subgenome-specific k-mers. New Phytologist, 235: 801–809.

Emms, DM, Kelly, S. 2019. OrthoFinder: phylogenetic orthology inference for comparative genomics. Genome Biology 20: 238.

Yang, Z. 2007. PAML 4: phylogenetic analysis by maximum likelihood. Molecular Biology and Evolution, 24: 1586–1591.

Xie, J, Chen, Y, Cai, G, Cai, R, Hu, Z, Wang, H. 2023. Tree Visualization By One Table (tvBOT): a web application for visualizing, modifying and annotating phylogenetic trees. Nucleic Acids Research, 51: W587–W592.

Sun, P, Jiao, B, Yang, Y, Shan, L, Li, T, Li, X, Xi, Z, Wang, X, Liu, J. 2022. WGDI: A user-friendly toolkit for evolutionary analyses of whole-genome duplications and ancestral karyotypes. Molecular Plant, 15: 1841–1851.

Buchfink, B, Reuter, K, Drost, H-G. 2021. Sensitive protein alignments at tree-of-life scale using DIAMOND. Nature Methods, 18: 366–368.

Edgar, RC. 2004. MUSCLE: multiple sequence alignment with high accuracy and high throughput. Nucleic Acids Research, 32: 1792–1797.

Suyama, M, Torrents, D, Bork, P. 2006. PAL2NAL: robust conversion of protein sequence alignments into the corresponding codon alignments. Nucleic Acids Research, 34: W609–W612.

Jaillon, O, Aury, JM, Noel, B, Policriti, A, Clepet, C, Casagrande, A, Choisne, N, Aubourg, S, Vitulo, N, Jubin, C et al. 2007. The grapevine genome sequence suggests ancestral hexaploidization in major angiosperm phyla. Nature, 449: 463–467.

Li, B, Dewey, CN. 2011. RSEM: accurate transcript quantification from RNA-Seq data with or without a reference genome. BMC Bioinformatics, 12: 323.

Wang, Y, Tang, H, Debarry, JD, Tan, X, Li, J, Wang, X, Lee, TH, Jin, H, Marler, B, Guo, H et al. 2012. MCScanX: a toolkit for detection and evolutionary analysis of gene synteny and collinearity. Nucleic Acids Research, 40: e49.

Chen, L, Hu, X, Zhao, S. 2002. Molecular research on Huyou origin. Acta Horticulturae Sinica, 29: 276–277.

Chen, S, Yang, H, Zheng, Y, Chen, Y, Qiu, Y. 2006. Preliminary study on identification of excellent genotypes of Changshan Huyou by molecular markers. Journal Of Molecular Cell Biology, 39: 502–508.

Xu, CJ, Bao, L, Zhang, B, Bei, ZM, Ye, XY, Zhang, SL, Chen, KS. 2006. Parentage analysis of huyou (*Citrus changshanensis*) based on internal transcribed spacer sequences. Plant Breeding, 125: 519–522.

Nakandala, U, Masouleh, AK, Smith, MW, Furtado, A, Mason, P, Constantin, L, Henry, RJ. 2023. Haplotype resolved chromosome level genome assembly of *Citrus australis* reveals disease resistance and other citrus specific genes. Horticulture Research, 10: uhad058.

Jang, MJ, Cho, HJ, Park, YS, Lee, HY, Bae, EK, Jung, S, Jin, HS, Woo, J, Park, E, Kim, SJ et al. 2024. Haplotype-resolved genome assembly and resequencing analysis provide insights into genome evolution and allelic imbalance in. Nature Genetics, 56: 2551–2561.

Djari, A, Madignier, G, Di Valentin, O, Gillet, T, Frasse, P, Djouhri, A, Hu, GJ, Julliard, S, Liu, MC, Zhang, Y et al. 2024. Haplotype-resolved genome assembly and implementation of VitExpress, an open interactive transcriptomic platform for grapevine. Proceedings of the National Academy of Sciences of the United States of America, 121: e2403750121.

Li, P, Xiao, L, Du, QZ, Quan, MY, Song, YP, He, YL, Huang, WX, Xie, JB, Lv, CF, Wang, D et al. 2023. Genomic insights into selection for heterozygous alleles and woody traits in. Plant Biotechnology Journal, 21: 2002–2018.

